# Barley MLA3 recognizes the host-specificity determinant PWL2 from rice blast (*M. oryzae*)

**DOI:** 10.1101/2022.10.21.512921

**Authors:** Helen J. Brabham, Diana Gómez De La Cruz, Vincent Were, Motoki Shimizu, Hiromasa Saitoh, Inmaculada Hernández-Pinzón, Phon Green, Jennifer Lorang, Koki Fujisaki, Kazuhiro Sato, István Molnár, Hana Šimková, Jaroslav Doležel, James Russell, Jodie Taylor, Matthew Smoker, Yogesh Kumar Gupta, Tom Wolpert, Nicholas J. Talbot, Ryohei Terauchi, Matthew J. Moscou

## Abstract

Plant nucleotide-binding leucine-rich repeat immune receptors (NLRs) directly or indirectly recognize pathogen-secreted effector molecules to initiate plant defense. Recognition of multiple pathogens by a single NLR is rare and usually occurs via monitoring for changes to host proteins; few characterized NLRs have been shown to recognize multiple effectors. The barley NLR *Mla* has undergone functional diversification and *Mla* alleles recognize host-adapted isolates of barley powdery mildew (*Blumeria graminis* f. sp. *hordei; Bgh*). Here, we show that *Mla3* also confers resistance to rice blast (*Magnaporthe oryzae*) in a dosage dependent manner. Using a forward genetic screen, we discovered that the recognized effector from *M. oryzae* is *PWL2*, a host range determinant factor that prevents *M. oryzae* from infecting weeping lovegrass (*Eragrostis curvula*). *Mla3* has therefore convergently evolved the capacity to recognize effectors from diverse pathogens.

## Introduction

Plants are routinely exposed to a diverse array of microbes and their interaction is governed by active processes that recognize self and non-self molecular patterns. The plant immune system is comprised of membrane-localized extracellular receptors and intracellular receptors that detect pathogen molecules or host-derived molecules generated during pathogen infection (Jones & Dangl, 2006). Plant pathogens routinely secrete effector molecules to promote virulence through manipulation of the host environment and suppression of the plant immune system. Effectors are highly sequence and structurally diverse molecules, evolving to evade plant recognition whilst also maintaining virulence function (Franceschetti *et al*., 2017). Nucleotide-binding, leucine-rich repeat (NLR) proteins are the largest class of immune receptors in plants and are grouped by their variable N-terminal domains: coiled coil (CC), Toll, interleukin-1 receptor, resistance protein (TIR), or RESISTANCE TO POWDERY MILDEW 8 (RPW8) domain (Ngou *et al*., 2022). The majority of characterized NLRs recognize single species- or isolate-specific effectors, initiating immune signaling and plant defense upon recognition (Kourelis & van der Hoorn, 2018).

The mechanism of recognition by NLRs is broadly classified by the direct and indirect perception of pathogen derived molecules (Kourelis & van der Hoorn, 2018; Saur *et al*., 2021). Direct recognition involves physical interaction of the NLR and effector to initiate NLR activation (direct model) (Jia *et al*., 2000), whereas indirect recognition by NLRs occurs through monitoring host targets for effector-mediated modifications (guard model) (Van Der Biezen & Jones, 1998). Host targets can also become integrated into NLRs within the same open reading frame via fusion events, forming additional domains for effector interaction and recognition (integration model) (Cesari, 2018; Cesari *et al*., 2014). These integrated NLRs routinely require a second NLR to initiate defense signaling (Cesari, 2018; Cesari *et al*., 2014). Extensive functional analysis has been performed on NLRs from each of these modes of recognition, however, the evolution and maintenance of these diverse recognition mechanisms is complex and often unclear (Märkle *et al*., 2022).

Recognition of effectors can be shared across plant species. Recognition is either conferred by related orthologous NLRs that perceive the same effectors or modified host targets (orthologous recognition), or by unrelated NLRs that have independently evolved a similar function (convergent recognition). The majority of examples of convergent recognition of pathogen effectors involve the guard or integration model of recognition. In *A. thaliana*, the NLR RPS5 recognizes AvrPphB through effector-mediated modification of the host serine/threonine protein kinase PBS1 (Ade *et al*., 2007; DeYoung *et al*., 2012; Qi *et al*., 2014; Shao *et al*., 2003). PBS1 is widely conserved in diverse plant species and proteolytic cleavage of homologs from *Arabidopsis*, barley (*Hordeum vulgare*), and wheat (*Triticum aestivum*) activates an immune response (Caldwell & Michelmore, 2009; Carter *et al*., 2019; Kim *et al*., 2016; Sun *et al*., 2017). The barley CC-NLR Pbr1 is unrelated to RPS5 yet also confers AvrPphB recognition, suggesting convergent recognition mediated through guarding of PBS1 (Carter *et al*., 2019). Another example of the guard model involves RIN4, a conserved component of the plant immune system. The *A. thaliana* NLRs RPM1 and RPS2 (Axtell & Staskawicz, 2003; Kim *et al*., 2005; Kim *et al*., 2009; Mackey *et al*., 2003; Mackey *et al*., 2002) and soybean (*Glycine max*) Rpg1b and Rpg1r (Ashfield *et al*., 2004; Ashfield *et al*., 2014; Russell *et al*., 2015) recognize RIN4 perturbation. All four NLRs are phylogenetically unrelated, thus multiple NLRs have convergently evolved the ability to monitor RIN4 (Toruño *et al*., 2019). In apple, proteolytic cleavage of RIN4 by AvrRpt2 from *Erwinia amylovora* also activates the NLR Mr5 (Prokchorchik *et al*., 2020). Tomato Ptr1 also recognizes AvrRpt2 via involvement of RIN4 homologs yet is unrelated to Mr5 or *A. thaliana* RPS2 (Mazo-Molina *et al*., 2020; Mazo-Molina *et al*., 2019). Across species, NLR activation occurs after different effector mediated modifications of RIN4 homologs (such as phosphorylation, acetylation, and proteolytic cleavage), supporting the hypothesis of multiple and convergent evolutionary origins (Kim *et al*., 2022).

Individual NLRs can interact with multiple guardees or host proteins to facilitate recognition of different effectors. The NLR ZAR1 from *A. thaliana* recognizes different bacterial effectors from *Xanthomonas campestris* and *Pseudomonas syringae* via individual interaction with the closely related pseudokinases PBL2, RKS1, ZED1, and ZRK3 (Lewis *et al*., 2013; Seto *et al*., 2017; Wang *et al*., 2015). The AtZAR1 signaling pathway is conserved in *Nicotiana benthamiana* and recognition of *X. perforans* effector XopJ4 is mediated by NbZAR1 and the receptor-like cytoplasmic kinase JIM2 (Baudin *et al*., 2017; Schultink *et al*., 2019). ZAR1 displays convergent recognition of diverse bacterial effectors through their interaction with different receptor-like cytoplasmic kinases across species.

Recognition of effectors from taxonomically diverse pathogens by the same NLR is hypothesized to have evolved through convergent processes. The paired NLRs RPS4/RRS1 confer resistance to the fungal pathogen *Colletotrichum higginsianum* and bacterial pathogens *Ralstonia solanacearum*, *Pseudomonas syringae*, and *Xanthomonas campestris* (Deslandes *et al*., 2003; Gassmann *et al*., 1999; Ma *et al*., 2018; Narusaka *et al*., 2017; Narusaka *et al*., 2013; Narusaka *et al*., 2009). Recognition occurs through perception of the effectors AvrRps4 from *P. syringae* and PopP2 from *R. solanacearum* with the integrated WRKY domain at the C-terminus of RRS1. AvrRps4 physically interacts with the WRKY domain, whereas recognition of PopP2 also requires its acetyl-transferase enzymatic activity on the integrated domain (Le Roux *et al*., 2015; Ma *et al*., 2018; Mukhi *et al*., 2021; Sarris *et al*., 2015; Saucet *et al*., 2015; Williams *et al*., 2014). In oat (*Avena sativa*), *Pc2* conditions resistance to *Puccinia coronata* (oat crown rust), yet also confers susceptibility to the host-selective toxin victorin from *Cochliobolus victoriae*. *Pc2* is uncharacterized in oat, but several additional plant species exhibit natural variation in victorin sensitivity including *A. thaliana* (Lorang *et al*., 2004), barley (Lorang *et al*., 2010), *Brachypodium* spp., and *Phaseolus vulgaris* (common bean) (Lorang *et al*., 2018). The NLR *AtLOV1* confers victorin sensitivity in *A. thaliana*, through the guarding of the thioredoxin *TRXh5* (Lorang *et al*., 2007; Sweat *et al*., 2008; Sweat & Wolpert, 2007). Victorin sensitivity in *P. vulgaris* is conferred by an unrelated NLR, although it is unknown whether a thioredoxin is involved in the interaction (Lorang *et al*., 2018). Therefore, several examples of convergent recognition have been observed in diverse angiosperm species, with the majority involving indirect recognition of a guarded host protein or via an integrated domain. Other NLRs that recognize taxonomically divergent pathogens include tomato *Mi-1* conferring resistance to root-knot nematodes (*Meloidogyne* spp.), potato aphid (*Macrosiphum euphorbiae*), and sweet potato whiteflies (*Bemisia* spp.) (Goggin *et al*., 2006; Nombela *et al*., 2003; Santos *et al*., 2020; Vos *et al*., 1998) and barley *Mla8* conferring resistance to barley powdery mildew (*Blumeria graminis* f. sp. *hordei*) and wheat stripe rust (*Puccinia striiformis* f. sp. *tritici*) (Bettgenhaeuser *et al*., 2021). *Mi-1* requires the partner NRC4 for cell death signaling (Wu *et al*., 2017), however *Mla8* does not contain an integrated domain and no interacting partners have yet been implicated for functional requirement. Across plant species, different NLRs have evolved to convergently recognize the same effectors or host targets highlighting conserved pathogen molecules or important components of the plant immune system. Conversely, NLRs have also evolved expanded recognition capabilities through the capacity to recognize sequence unrelated effectors.

The barley *Mla* locus on the short arm of chromosome 1H is a resistance gene complex showing extreme intraspecific diversity in barley haplotypes (Briggs & Stanford, 1938; Jørgensen & Wolfe, 1994; Seeholzer *et al*., 2010; Wei *et al*., 2002; Zhou *et al*., 2001). Characterization of the *Mla* locus in the reference genome Morex identified a complex region containing three NLR gene families—*RGH1 (Mla)*, *RGH2*, and *RGH3*—that are located in three gene-rich regions flanked by repetitive and mobile elements (Wei *et al*., 2002). Allelic variants of the *Mla* CC-NLR gene (*RGH1*) confer isolate-specific immunity against the host pathogen barley powdery mildew *Bgh* (Halterman *et al*., 2001; Jørgensen & Wolfe, 1994; Kinizios *et al*., 1995; Seeholzer *et al*., 2010). The LRR region of *Mla* determines recognition specificity and shows signatures of positive selection (Maekawa *et al*., 2019; Seeholzer *et al*., 2010; Shen *et al*., 2003). A direct recognition mechanism for *Mla* has been proposed, shown by the direct interaction between MLA1, MLA7, MLA10, MLA13, and MLA22 and the *Bgh* effectors AVR_a1,_ AVR_a7_, AVR_a10_, AVR_a13_, and AVR_a22,_ respectively (Bauer *et al*., 2021; Lu *et al*., 2016; Saur *et al*., 2019).

Recognition of diverse pathogens has been genetically linked to the *Mla* locus including susceptibility to *Bipolaris sorokiniana* (*Rcs6*) (Bilgic *et al*., 2006; Leng *et al*., 2020; Leng *et al*., 2018) and resistance to *Magnaporthe oryzae* (syn. *Pyricularia oryzae*; *Rmo1*) (Inukai *et al*., 2006). Previous work found that *Resistance to* M. oryzae *1* (*Rmo1*) was in genetic coupling with the *Mla* locus in the barley accession Baronesse (*Mla3*) (Inukai *et al*., 2006). In this study, we perform a high-resolution recombination screen and find that *Rmo1* is in complete genetic coupling with *Mla3*. Using RNAseq, RenSeq-PacBio, and chromosome sequencing, we show that all three NLR gene families at the *Mla3* locus—*RGH1*, *RGH2*, and *RGH3*—are present and expressed in Baronesse. Using whole genome sequencing and *k*-mer analysis, we report that Baronesse carries four near-identical copies of *Mla3*. Characterization of a diversity panel and *Mla* introgression lines suggested that *Mla3* underlies *Rmo1*-mediated resistance. *Agrobacterium*-mediated transformation provides evidence that *Mla3* specifically conferred resistance to *M. oryzae* isolate KEN54-20 (*AVR-Rmo1*) in a dosage-dependent manner. Using association genetics and mutagenesis combined with high-throughput sequencing, we report the unexpected finding that *Mla3* (*Rmo1*) recognizes a known host species specificity determinant *PWL2* (*AVR-Rmo1*).

## Results

### *Mla3* is in complete genetic coupling with *Rmo1*

To elucidate the genetic relationship of *Mla3* and *Rmo1*, a recombination screen using a Baronesse x BCD47 F_2_ population (N=2,304 gametes) was performed over a wide 22.9 cM region on chromosome 1H encompassing the *Mla* locus (markers K_3_1144 and K_2_0712) (**Figure 1a**). Among 169 recombinants, 80 recombinants were identified between the flanking markers of the *Mla* locus (K_963924 and K_206D11). Marker saturation of the region encompassing *Mla* was performed using 12 KASP markers spanning a wide genetic interval from markers K_3_0933 to K_4261. Homozygous recombinants were identified for 53 F_2:3_ families. Twenty-four additional KASP markers were generated by identifying single nucleotide polymorphisms (SNPs) between Baronesse and BCD47 based on PCR amplification of the *Mla* locus. Suppressed recombination was observed at the *Mla3* locus, with 18 markers in complete genetic coupling with K_Mla_RGH1, that spans a ~240 kb physical region in the reference sequence cv. Morex [Wei 2002]. The marker K_Mla_RGH1 was designed on a SNP between *Mla3* coding sequence and the *RGH1* allele in BCD47 (*RGH1-BCD47*). A total of 165 F_2:3_ families with recombination events within the interval surrounding the *Mla3* locus were phenotyped with *Bgh* isolate CC148 (*AVR_a3_*), to determine if the marker K_Mla_RGH1 is in coupling with barley powdery mildew (*Bgh*) resistance. Using these 165 F_2:3_ families, resistance to *Bgh* isolate CC148 cosegregated with marker K_Mla_RGH1. Baronesse and BCD47 show clear differential phenotypes upon inoculation with the *M. oryzae* isolate KEN54-20 carrying *AVR-Rmo1*. To map *Rmo1*, 42 homozygous F_2:4_ families with recombination events near the *Mla* locus were inoculated with *M. oryzae* isolate KEN54-20 using leaf-spot inoculation (**Figure 1b**) and whole plant spray inoculation (**Figure 1c**). Resistance to *M. oryzae* isolate KEN54-20 cosegregated with marker K_Mla_RGH1. Therefore, *Rmo1* is in complete genetic coupling with *Mla3*.

**Figure 1.**
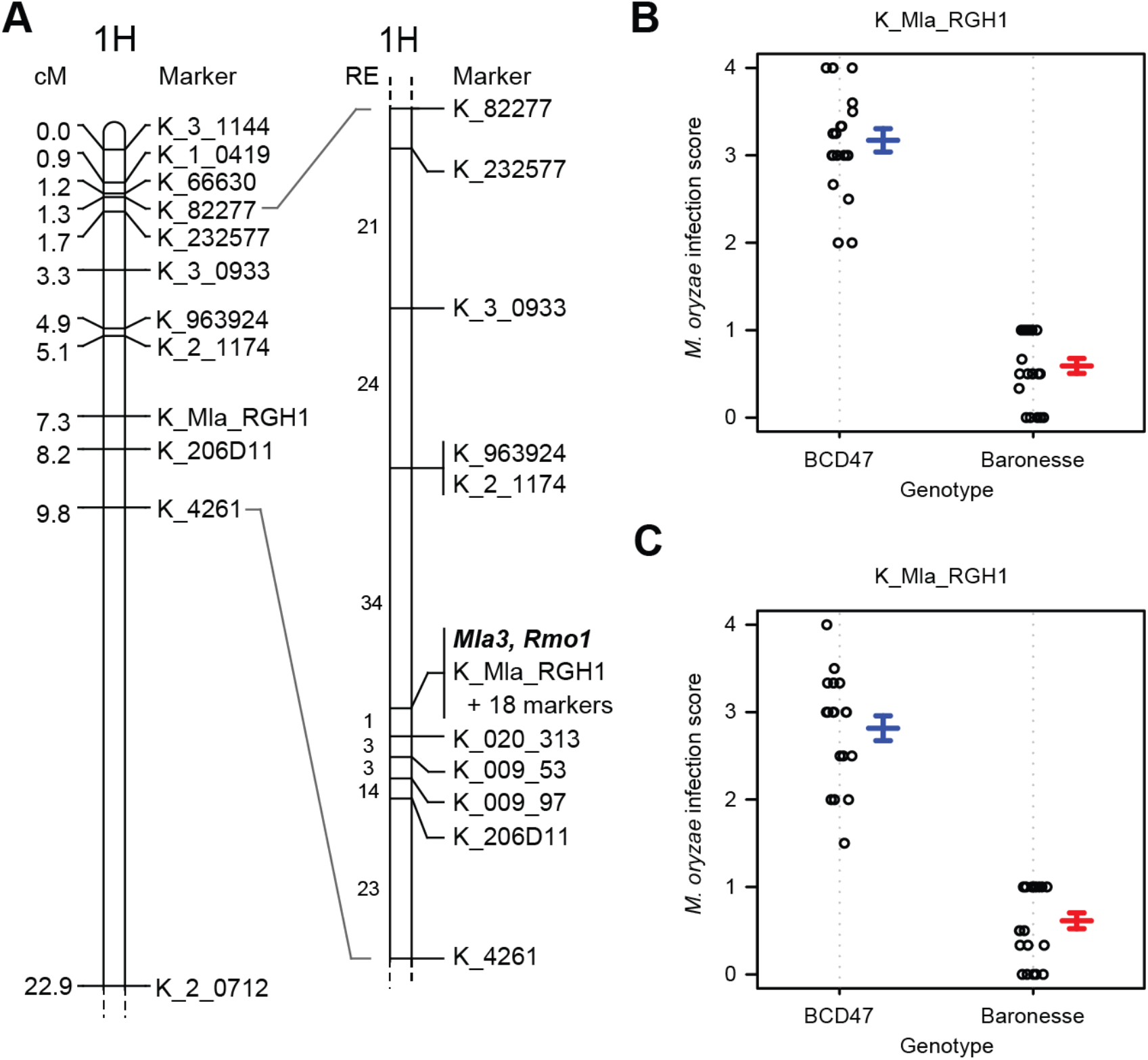
*Rmo1* is in complete coupling with *Mla3*. **(A)** The distal end of the short arm of chromosome 1H based on non-redundant KASP markers in the Baronesse x BCD47 population (2,304 gametes). RE, number of recombination events between markers. Twenty additional markers (not shown) are in complete genetic coupling with K_Mla_RGH1_2920 at the *Mla* locus. **(B)** Phenotype by genotype plot of homozygous F_2:4_ recombinants inoculated with *M. oryzae* isolate KEN54-20 using a spot-based inoculation. Scores 0 and 1 = resistant and 2 to 4 = susceptible. **(C)** Phenotype by genotype plot of homozygous F_2:4_ recombinants inoculated with *M. oryzae* isolate KEN54-20 using a spray-based inoculation. Scores 0 and 1 = resistant and 2 to 4 = susceptible.

### *Mla3* (*RGH1*), *RGH2*, and *RGH3* are present and expressed in the *Mla3* haplotype from Baronesse

The physical size, structure and gene content of the *Mla* locus in Baronesse is unknown. The *Mla* locus in the reference genome of Morex was first sequenced using a bacterial artificial chromosome approach (Wei *et al*., 1999; Wei *et al*., 2002). The locus fails to assemble correctly using short read sequencing technology due to high repetitive content and presence of large near-identical duplications (Mascher *et al*., 2021). In Morex, the *Mla* locus includes multiple members of three CC-NLR gene families—*RGH1*, *RGH2*, and *RGH3*—of which all three are present within a 40 kb tandem duplication (Wei *et al*., 2002). We used several approaches to resolve the genomic region encompassing *Mla3*. Using chromosome flow sorting and Chicago long-range linkage (Thind *et al*., 2017), we assembled chromosome 1H from barley accession Baronesse. Genomic scaffolds were fragmented at the boundaries of the *Mla* locus, with the distal and proximal regions defined by scaffold1235 and scaffold1874, respectively (**Figure 2a; Supplemental Figure 1**). Genomic sequence of *Mla3* (*RGH1*) was fragmented, with fragments found on scaffold1874 and three small contigs (contigs 38297, 42637, and 63307), whereas *RGH2* and *RGH3* were found to be in a head-to-head orientation on scaffold1874. The flanking intervals for the *Mla3* haplotype are highly co-linear with the Morex haplotype (**Figure 2a; Supplemental Figure 1**). Using a complementary approach, we applied RenSeq-PacBio (Witek *et al*., 2016) with a capture library designed on NLRs that included baits designed on *Mla3* (*RGH1*), *RGH2*, and *RGH3* gene families (Brabham *et al*., 2018). *Mla3* and *RGH2*/*RGH3* were assembled on 15.3 and 14.0 kb contigs, respectively (**Figure 2b**).

**Figure 2.**
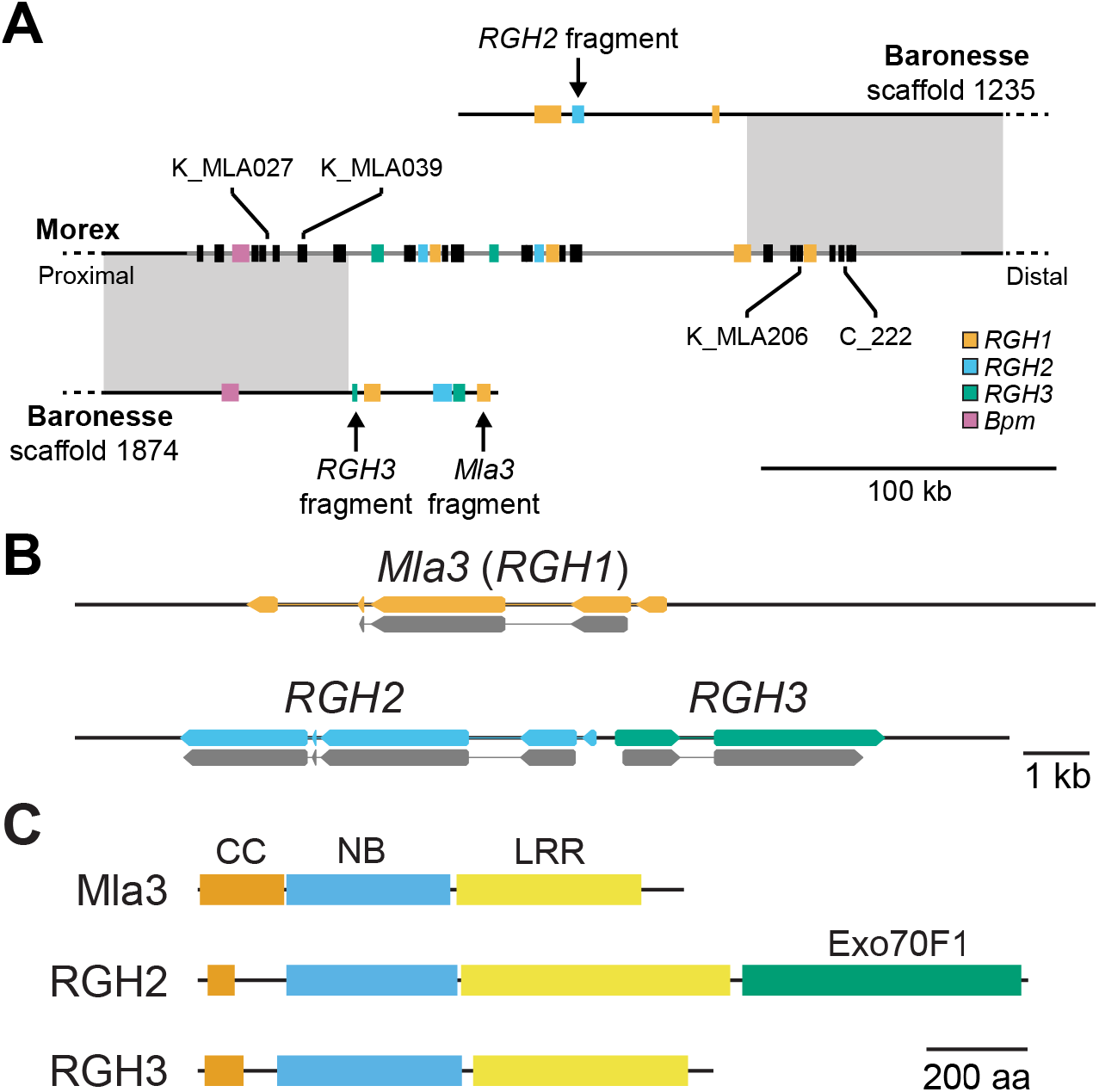
The *Mla* locus in Baronesse is highly divergent in sequence and structure. **(A)** Sequence alignment of the region encompassing *Mla* in barley accessions Morex (*mla*) and Baronesse (*Mla3*) found high conservation in the flanking intervals (grey boxes) but no conservation within the *Mla* locus. The central region of *Mla* region is a breakpoint in the assembly of chromosome 1H from barley accession Baronesse. *RGH1*, *RGH2*, and *RGH3* family members are indicated in orange, blue, and green, respectively. KASP and CAPS markers are indicated with ‘K_’ and ‘C_’ prefix, respectively. **(B)** RenSeq-PacBio identified genomic contigs encompassing *Mla3* (15.3 kb) and *RGH2*/*RGH3* (14.0 kb). In the barley accession Baronesse haplotype, *RGH2* and *RGH3* are in head-to-head orientation. **(C)** In the barley accession Baronesse haplotype, Mla3 and RGH3 encode CC-NB-LRR, whereas RGH2 encodes an CC-NB-LRR with integrated Exo70F1.

We hypothesized that the fragmented assembly is due to repetitive gene content or large near-identical duplications, such as observed in the reference genome (Mascher *et al*., 2021). To identify potential duplications, we used *k*-mer counting with flow sorted chromosome 1H sequencing data using genomic contigs encompassing *Mla3* and *RGH2*/*RGH3* identified by RenSeq-PacBio as template (**Figure 3**). Based on local regression of *k*-mer coverage, the single copy gene *Bpm* located immediately proximal to the *Mla* locus had an estimated coverage of 161.6 (5’ region) and 165.1 (3’ region) (**Figure 3a**), whereas *Mla3* had an estimated coverage of 800.1 (**Figure 3b**). Evaluation of individual *k*-mer coverage found that the average coverage is an overestimate due to high-copy *k*-mers present in the central region of *Mla3*. Analysis using *k*-medoids found that individual *k*-mers of *Mla3* clustered at 146, 619, and 790, corresponding to one, four, and five copies, respectively (**Supplemental Figure 2**). The same approach found *Bpm* and *RGH2/RGH3* had *k*-medoids of 175 (k=1) and 162 and 315 (k=2), respectively (**Supplemental Figure 2**). *RGH2* and *RGH3* are single copy, although fragments of the C-terminal encoding regions of the NLRs are present in scaffolds 1235 and 1874, respectively, that flank the *Mla* locus (**Figure 2a**, **Figure 3c**). For *Mla3*, *k*-mers contributing to five copies are concentrated in the regions encoding the coiled coil and nucleotide binding domains (**Figure 3b**). Two regions in the leucine-rich repeat region of *Mla3* were found to have a reduction in *k*-mer coverage suggesting that a single copy of *Mla3* had diverged from other copies (**Figure 3b**). Evaluation of aligned genomic reads found heterogeneity in *Mla3* copies, with one copy carrying a six base pair deletion resulting in two deletions and one change in the amino acid sequence. This copy was designated *Mla3Δ6*. Collectively, based on this analysis we conclude that Baronesse carries three identical copies of *Mla3*, *Mla3Δ6*, and an *RGH1* family member that is diverged from *Mla3*.

**Figure 3.**
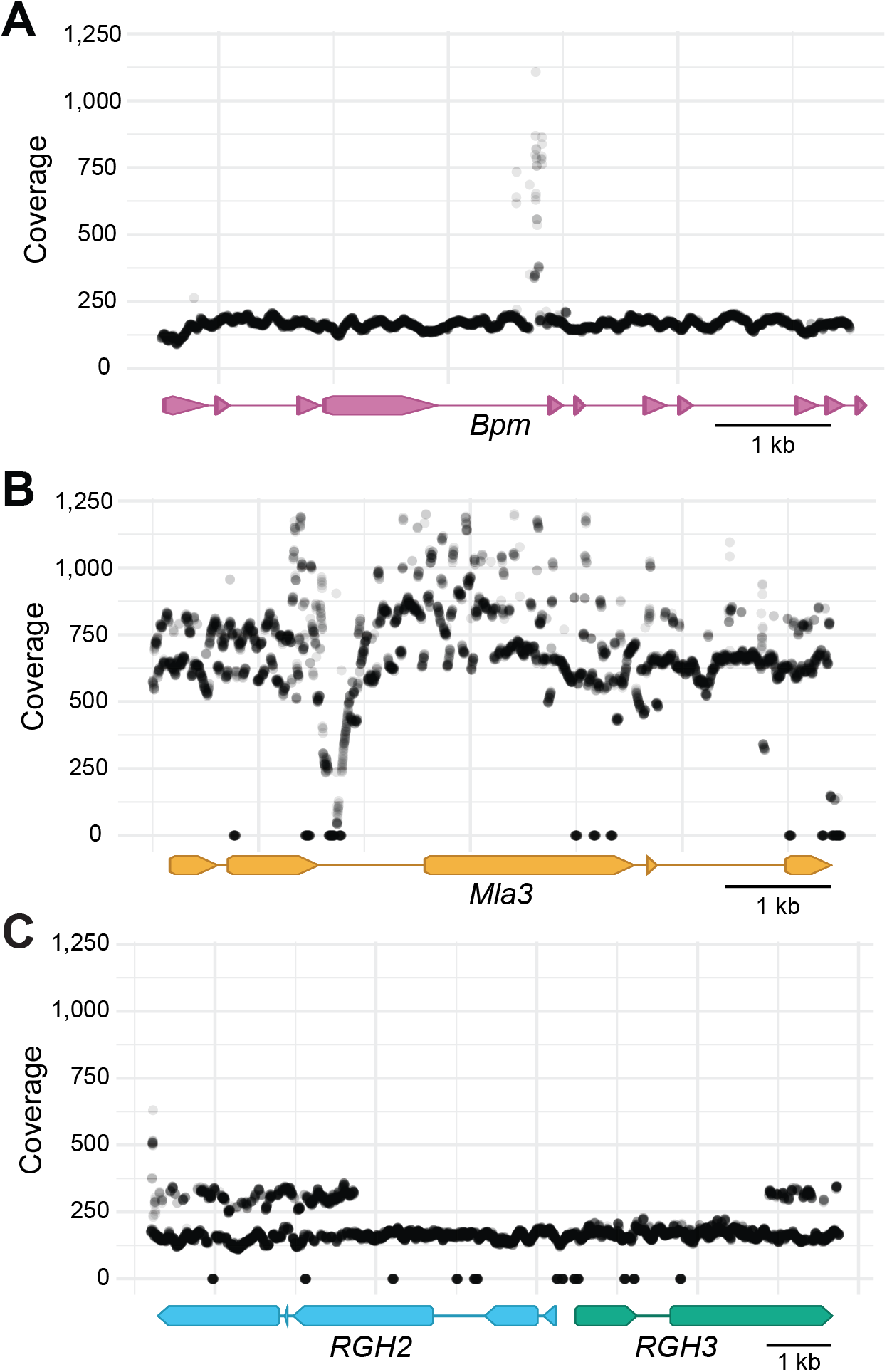
*k*-mer analysis identifies four copies of *Mla3* in barley accession Baronesse. **(A)** *k*-mer coverage of *Bpm*. *Bpm* is located proximal to the *Mla* locus and exists as a single copy with coverage centered at 175. **(B)** *k*-mer coverage of *Mla3*. Two coverage bands are observed at 619 and 790 coverage, corresponding to four and five copies. Reduced coverage in the second intron is due to the presence of low complexity sequence (dinucleotide repeat). **(C)** *k*-mer coverage of *RGH2* and *RGH3*. Two coverage bands are observed at 162 and 315 coverage, corresponding to one and two copies, respectively. The additional copies represent fragments of *RGH2* and *RGH3*, which are located in the distal and proximal boundaries of the *Mla* locus (**Figure 2a**). Zero *k*-mer coverage represents inaccurate sequence calls from RenSeq-PacBio sequencing in contigs encompassing *Mla3* and *RGH2*/*RGH3*. Gene models are shown below each plot with exons and introns shown as arrows and lines, respectively.

*De novo* transcriptome assembly of RNAseq from first leaf of Baronesse found that *Mla3*, *RGH2*, and *RGH3* family members were expressed. To confirm expression of the *Mla3Δ6* copy and identify other expressed variants, we aligned RNAseq onto gene models of *Mla3*, *RGH2*, and *RGH3*. No variation was found in *RGH2* and *RGH3*, whereas we verified the existence two expressed variants of *Mla3* (*Mla3* and *Mla3Δ6*). Baronesse *RGH2* encodes an NLR with an integrated Exo70F1 and is in head-to-head orientation with *RGH3* (**Figure 2c**). *RGH2* and *RGH3* belong to the Major Integration Clade 1 (MIC1) and C7 clades of NLRs, respectively (Bailey *et al*., 2018; Brabham *et al*., 2018). Other members of the MIC1 clade include *Rpg5* from barley (Wang *et al*., 2013) and *RGA5* from rice (Cesari *et al*., 2014), which require additional NLRs to function as a pair. Their respective partners, *RGA1* (Wang *et al*., 2013) and *RGA4* (Cesari *et al*., 2014) also reside in the C7 clade. Following this observation, we hypothesize that *RGH2* and *RGH3* also function as paired NLRs. Therefore, candidate genes for conferring *Rmo1* resistance are *Mla3*, *Mla3Δ6*, and the paired NLRs *RGH2* and *RGH3*.

### Baronesse *RGH2* and *RGH3* do not confer *Rmo1-*mediated resistance

An Exo70 in rice, OsExo70F3, is the target of the *M. oryzae* effector AVR-Pii and this interaction is guarded by the resistance gene pair *Pii* and *Pii-2* (Fujisaki *et al*., 2015). We hypothesized that Exo70s could be a conserved effector target between rice and barley, hence *RGH2* and *RGH3* are candidates for conferring *Rmo1*-mediated resistance. Barley haplotypes contain extensive variation at the *Mla* locus (Brabham *et al*., 2018; Maekawa *et al*., 2019; Seeholzer *et al*., 2010). To leverage this natural variation, we investigated *RGH2* and *RGH3* in a panel of over 40 diverse barley accessions *de novo* assembled transcriptomes (**Supplemental Data 1**) (Brabham *et al*., 2018; Maekawa *et al*., 2019). We identified multiple accessions containing diverse allelic variants of *RGH2* and *RGH3*. Of these, the accession Maritime contains identical copies of *RGH2* and *RGH3* as found in the accession Baronesse but carries a divergent *Mla* allele. Screening the diversity panel with *M. oryzae* isolate KEN54-20 (*AVR-Rmo1*) using both a spray- and spot-based inoculation found that all accessions, aside from Baronesse, were susceptible (**Figure 4; Supplemental Figure 3**). Therefore, we conclude that *RGH2* and *RGH3* do not confer *Rmo1*-mediated resistance.

**Figure 4.**
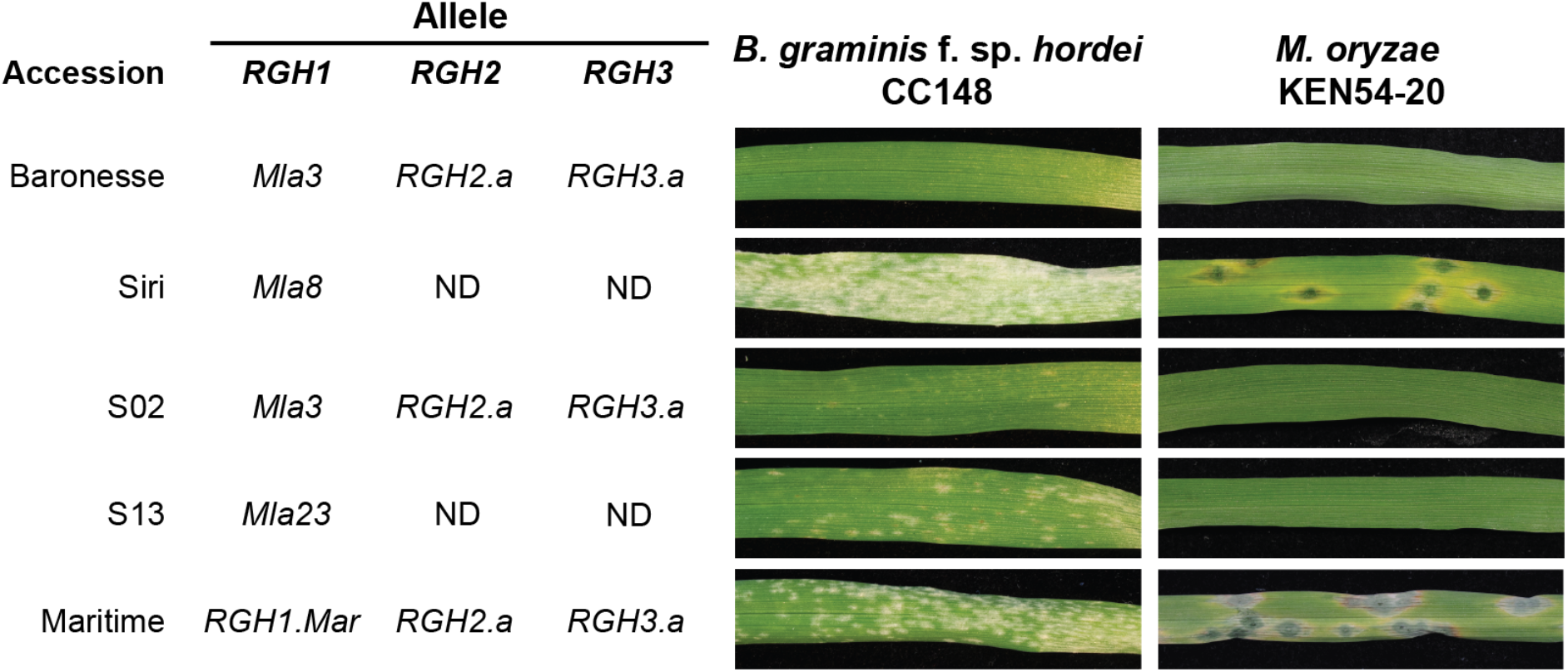
Coupling of *Mla3* and *Rmo1* in diverse barley accessions. Disease phenotypes of barley accessions carrying different alleles of the candidate genes *RGH1*, *RGH2*, and *RGH3* inoculated with *Bgh* isolate CC148 and *M. oryzae* isolate KEN54-20. The haplotype of the barley accessions are listed on the *left-hand* side with the allele of *RGH1*, *RGH2*, and *RGH3* indicated. ND, gene was not detected in RNAseq data.

### *Mla3* is *Rmo1*

To assess *RGH1* candidacy for *Rmo1* resistance, we took advantage of an introgression panel containing diverse mildew resistance loci. The Siri near-isogenic lines (NILs) were generated by crossing donor accessions with the recurrent parent Siri (*Mla8*), followed by multiple rounds of backcrossing and selection using *Bgh* isolates (Kølster & Stølen, 1987). The Siri NILs contains 13 mildew resistance genes, including 11 *Mla* specificities, in isogenic background of Siri including *Mla1*, *Mla3*, *Mla6*, *Mla7* (Nordal), *Mla7* (Moseman), *Mla9*, *Mla10*, *Mla12*, *Mla13*, *Ml22*, *Mla23*, *Ml-(Ru2)*, and *Mlk*. Inoculation of the Siri NILs with *M. oryzae* isolate KEN54-20 using a spray- and spot-based inoculation found that line S02 containing *Mla3* and the line S13 carrying *Mla23* were resistant to KEN54-20 (**Figure 4; Supplemental Figure 4**). *Mla23* is the most closely related *Mla* allele to *Mla3*, sharing 98% sequence similarity at the DNA and protein level, with variation limited to the C-terminal region of the LRR (**Supplemental Figure 5**) (Seeholzer *et al*., 2010). Leaf RNAseq data of S13 confirmed the expression of Mla23 but did not detect *RGH2* or *RGH3*. The close phylogenetic relationship between *Mla3* and *Mla23* supports the hypothesis that *Mla3* underlies *Rmo1*-mediated resistance.

To determine whether *Mla3* or other NLRs present at the *Mla3* locus confer *Rmo1*-mediated resistance, all four candidate genes were cloned via PCR amplification—*Mla3* and *Mla3Δ6* from cDNA and *RGH2* and *RGH3* from gDNA. *Mla3* and *Mla3Δ6* were placed in an expression construct containing the *Mla6* promoter and terminator (**Supplemental Figure 6**). The *Mla6* promoter/terminator system was selected for direct comparison of *Mla3* and *Mla3Δ6* by eliminating native promoter variation. *RGH2* and *RGH3* were maintained in the native form and head-to-head orientation (**Supplemental Figure 6**). All constructs were transformed into the barley accession Golden Promise via *Agrobacterium* mediated transformation of immature embryos (Hensel & Kumlehn, 2004). Golden Promise is susceptible to *Bgh* isolate CC148 (*AVR_a3_*) and *M. oryzae* isolate KEN54-20 (*AVR-Rmo1*). *Mla3*, *Mla3Δ6*, and *RGH2*/*RGH3* transgenic Golden Promise T_1_ families were tested with the *Bgh* isolate CC148 (*AVR_a1_*, *AVR_a3_, avr_a6_*). Eight seed from two spikes were evaluated per family. Four independent T_1_ families were evaluated for *Mla3* (T_1_-3, T_1_-4, T_1_-5, and T_1_-6); four T_1_ families were evaluated for *Mla3Δ6* (T_1_-2, T_1_-3, T_1_-6, and T_1_-7); and seven T_1_ families for *RGH2*/*RGH3* (T_1_-1, T_1_-3, T_1_-4, T_1_-5, T_1_-9, T_1_-11, and T_1_-12). Only full-length *Mla3* was shown to confer resistance to *Bgh* isolate CC148, with all transgenic *Mla3Δ6* lines displaying susceptibility (**Figure 5a**). *Mla3Δ6* is 98% identical in coding and protein sequence to *Mla3*. *Mla3Δ6* could represent a loss-of-function allele or pseudogene of *Mla3*. All *RGH2*/*RGH3* transgenic barley T_1_ families were susceptible to *Bgh* isolate CC148 (**Figure 5a**).

**Figure 5.**
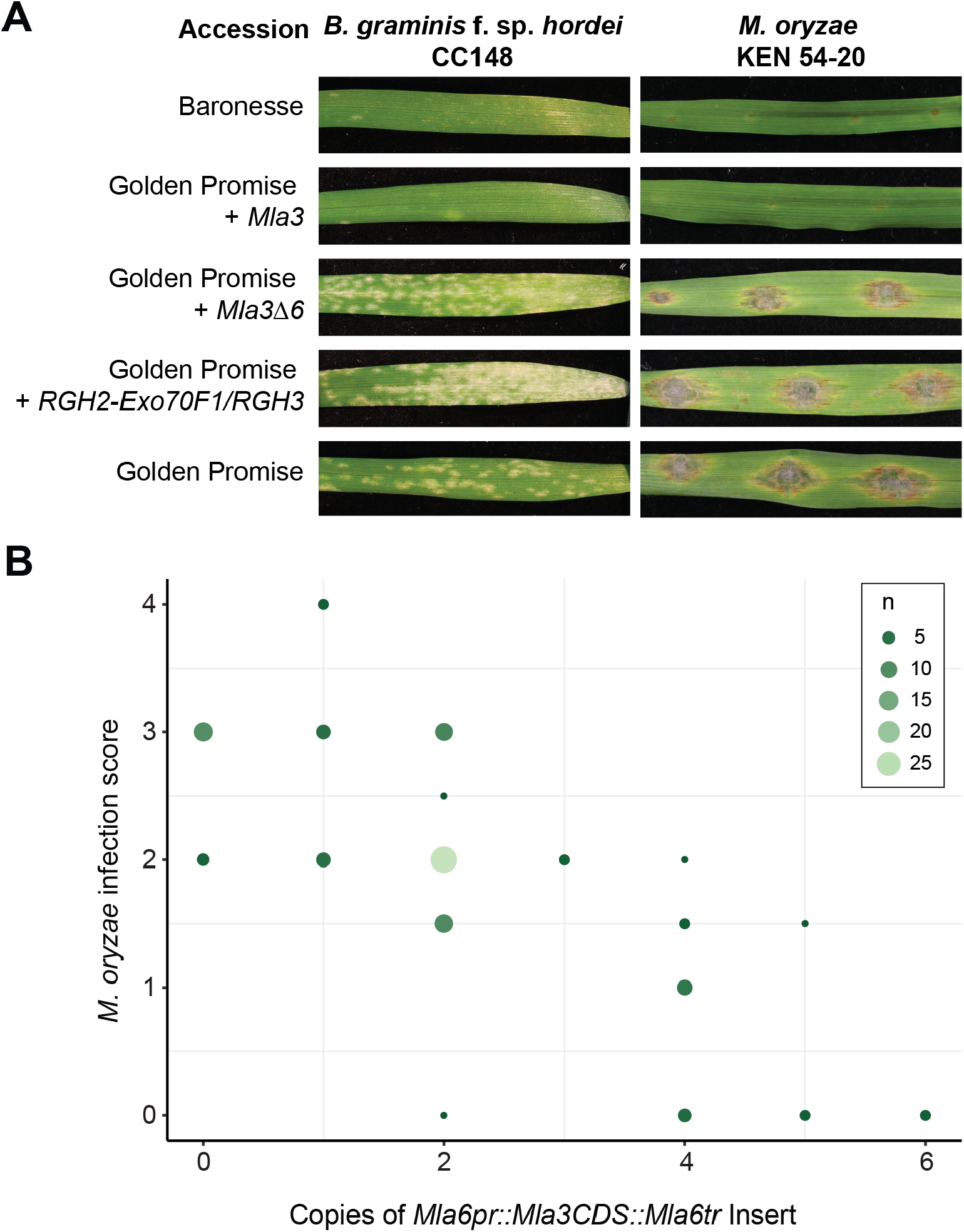
*Mla3* confers resistance to *Bgh* isolate CC148 and *M. oryae* isolate KEN54-20. **(A)** Transgenic lines of *Mla3* (T_1_-4), *Mla3Δ6* (T_1_-7), and *RGH2*/*RGH3* (T_1_-9) inoculated with *Bgh* isolate CC148 carrying *AVR_a3_,* and transgenic lines of *Mla3* (T_1_-4), *Mla3Δ6* (T_1_-3 T_2_), and *RGH2*/*RGH3* (T_1_-5) *M. oryzae* isolate KEN54-20. Controls include resistant wild-type Baronesse and susceptible wild-type Golden Promise used for transformation. Complete resistance shown to *Bgh* and *M. oryzae* by Baronesse and transgenic line Golden Promise + *Mla3*, whereas wild-type Golden Promise and transgenic lines Golden Promise + *Mla3Δ6* and + *RGH2/RGH3* are susceptible. Phenotypes are representative of inoculated T_1_ families. **(B)** Two independent T_1_ families of Golden Promise + *Mla3* (T_1_-4 and T_1_-5) showing resistance with varying copy number (0 to 6) inoculated with *M. oryzae* isolate KEN54-20. Phenotypic scores 0 and 1 = resistant; 2 to 4 = susceptible. Number of individual lines from T_1_ families with insert copy number for each phenotypic score, circle size and color gradient indicate number of individuals at each plot point (small dark green circles <=5 through to large light green circles = 25).

*Rmo1* confers dominant, race-specific resistance to *M. oryzae* isolate KEN54-20 (*AVR-Rmo1*) (Inukai *et al*., 2006). *Mla3*, *Mla3Δ6*, and *RGH2*/*RGH3* transgenic T_1_ families were screened with *M. oryzae* isolate KEN54-20 using spot inoculation on detached leaves. Eight seeds from two spikes were evaluated per family. Four independent T_1_ families were evaluated for *Mla3* (T_1_-3, T_1_-4, T_1_-5, and T_1_-6); four T_1_ families were evaluated for *Mla3Δ6* (T_1_-2, T_1_-3, T_1_-6, and T_1_-7); and seven T_1_ families for *RGH2*/*RGH3* (T_1_-1, T_1_-3, T_1_-4, T_1_-5, T_1_-9, T_1_-11, and T_1_-12). *Mla3* transgenic lines showed resistance to *M. oryzae* KEN54-20, recapitulating the wild-type phenotype (**Figure 5a; Supplemental Data 2**). Analysis of segregating *Mla3* transgenic T_1_ families showed phenotypic variation, with some families displaying partial or no resistance. In contrast, *Mla3Δ6* transgenic lines were susceptible to KEN54-20. All *RGH2*/*RGH3* transgenic individuals were fully susceptible. Collectively, these data show that resistance to both *Bgh* and *M. oryzae* was only observed in transgenic families carrying *Mla3*, although substantial intrafamily variation was observed for *M. oryzae*.

### Multiple copies of *Mla3* are required for *M. oryzae* resistance

Based on the observation of variable expression in *Mla3* T_1_ families for resistance to *M. oryzae* KEN54-20, we hypothesized that sufficient expression of *Mla3* is required to confer resistance. Natively, copy number variation is observed in wild-type Baronesse with three copies of *Mla3* and one copy of *Mla3Δ6* in the haploid genome. Therefore, we evaluated copy number variation of the individual transgenic lines. For the *Mla3* T_1_ families (T_1_-1, T_1_-4, T_1_-5, and T_1_-6) the number of copies ranged from 0 to 6. For the *Mla3Δ6* T_1_ lines, the number of copies of the transgenic insert varied from 1 to 4. Copy number analysis of the *RGH2/RGH3* T_1_ lines varied from 0 to 4 copies. Evaluation of copy number variation in T_1_ families and their phenotypic response to *M. oryzae* isolate KEN54-20 found an inverse linear correlation (**Figure 5b**). Multiple copies of *Mla3* were required to complement the wild-type phenotype and confer complete resistance to *M. oryzae* isolate KEN54-20 (**Figure 5b**). Resistance was only observed in individuals carrying greater than two copies of the insert, with a single copy being insufficient for complementation (**Figure 5b**).

Due to the high copy number being required for complementation in the transgenic lines, it was unclear if the observed resistance could be due to auto-activity of the transgene. Overexpression of NLRs has been shown to cause constitutive defense activation and broad-spectrum disease resistance to multiple pathogens (Lai & Eulgem, 2018; Li *et al*., 2019). To evaluate this, resistant *Mla3* transgenic lines were tested with *M. oryzae* isolate Sasa2 (*avr-Rmo1*), which is virulent on wild-type Baronesse carrying *Mla3*. Transgenic *Mla3* lines with high copy number that previously displayed resistance to *M. oryzae* isolate KEN54-20 were spot inoculated with *M. oryzae* isolates KEN54-20 and Sasa2 in single spots in proximal and distal positions on detached leaves (**Figure 6**). *Mla3* shows specific recognition of *M. oryzae* isolate KEN54-20 similar to wild-type Baronesse, but was susceptible to *M. oryzae* isolate Sasa2, showing large susceptible lesions (**Figure 6**). The barley accessions Golden Promise and Nigrate are susceptible to *M. oryzae* isolates KEN54-20 and Sasa2. Resistance to *M. oryzae* isolate KEN54-20 was maintained regardless of spot inoculation position on the leaf. Thus, *Mla3* provides isolate-specific resistance to KEN54-20 hypothesized through recognition of the effector *AVR-Rmo1* present in KEN54-20 but not Sasa2.

**Figure 6.**
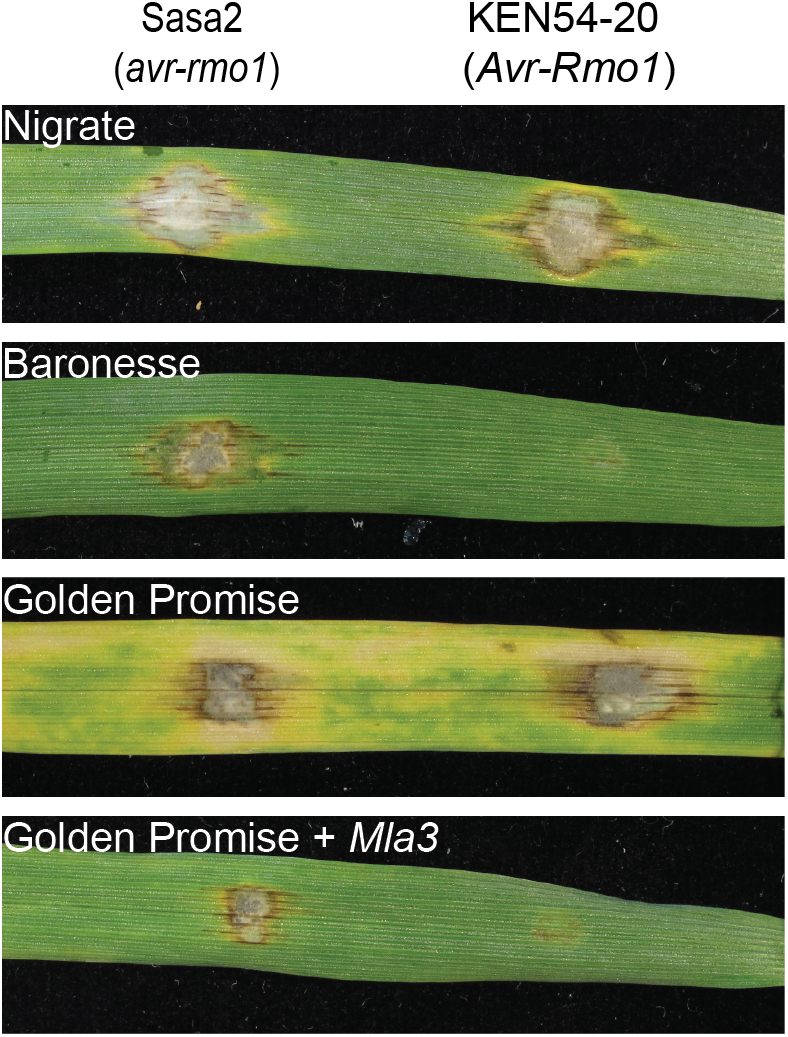
*Mla3* confers isolate-specific resistance to *M. oryzae*. Transgenic Golden Promise + *Mla3* (T_1_-4 T_2_) spot inoculated with *M. oryzae* isolates KEN54-20 (*AVR-Rmo1*) and Sasa2 (*avr-Rmo1*). *Mla3* confers resistance to isolates carrying *AVR-Rmo1*. Controls Golden Promise and hyper-susceptible Nigrate are susceptible to both Sasa2 and KEN54-20.

### *AVR-Rmo1* is *Pwl2*

To identify the effector *AVR-Rmo1* that is recognized by *Mla3*, we performed mutagenesis on spores of *M. oryzae* isolate KEN54-20 through exposure to UV light and screened for gain-of-virulence mutants on wild-type Baronesse following spray-based inoculation. A total of 72 putative mutants were identified in this primary screen. Loss of virulence (*avr-rmo1*) was confirmed in twelve mutants following re-inoculation on Baronesse using a spot-based method on both wild-type Baronesse (*Mla3*) and *Mla3* transgenic lines (T_1_-4 T_2_) (**Supplemental Figure 7**). To identify *AVR-Rmo1*, we first sequenced the genome of *M. oryzae* isolate KEN54-20 using the hybrid assembler MaSuRCA that uses long-read (Oxford Nanopore Technologies) and short-read (Illumina) technologies (Zimin *et al*., 2013). The final assembly was 47.9 Mb on 39 contigs. To identify shared mutations in the mutants, we performed whole genome sequencing and compared to the wild-type KEN54-20 genome. A window-based *k*-mer analysis was used to identify regions harboring SNPs, insertions, or deletions in all mutant lines. A region of 8 kb encompassing the known effector *PWL2* was deleted across all mutant lines (**Supplemental Figure 8**) (Sweigard *et al*., 1995). Due to repetitive regions in the flanking sequence of *PWL2*, the exact boundaries of the deletions in each mutant are ambiguous. Manual inspection of aligned reads to the region confirmed the deletion of the *PWL2* coding sequence in each mutant line. Therefore, the effector *PWL2* is a candidate for *AVR-Rmo1*. Gain of virulence to *Mla3* could also be due to other independent mutations in each mutant line or loss of a closely linked gene in the shared *PWL2* deletion. To determine whether *PWL2* is *AVR-Rmo1,* we complemented mutant *M. oryzae* lines with *PWL2* expressed under its native promoter. Independent ectopic transformed mutant lines of *M. oryzae* (M43*+pPWL2:PWL2* transformant 4 and M61*+pPWL2:PWL2* transformant 1) were avirulent on Baronesse (*Mla3*) and transgenic *Mla3* plants (**Figure 7ab)** confirming that *PWL2* is *AVR-Rmo1*.

**Figure 7.**
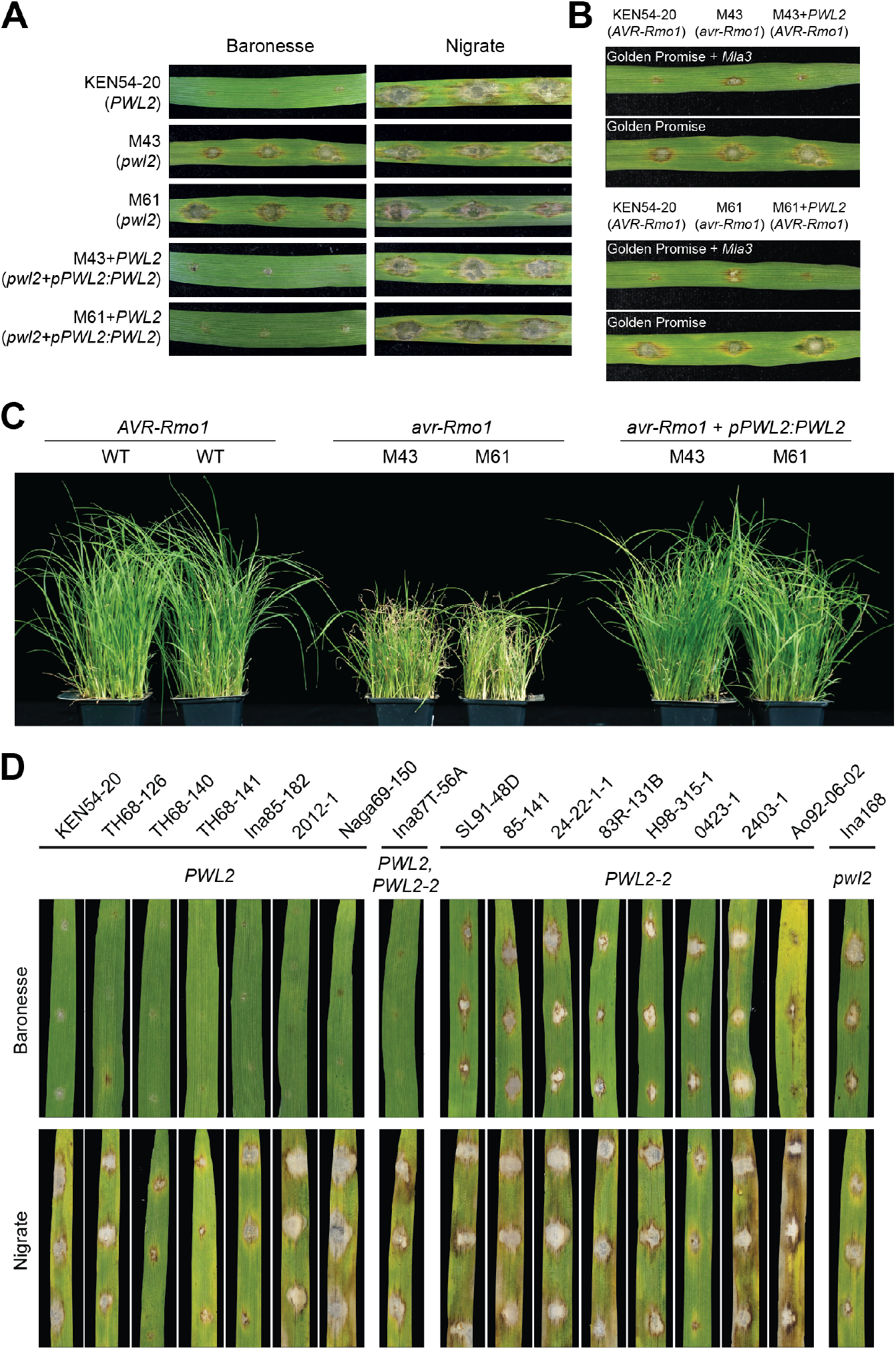
Conserved recognition specificity of *PWL2* in barley and weeping lovegrass. **(A)** *PWL2* complements *M. oryzae avr-Rmo1* mutants. Baronesse (*Mla3*) leaves spot inoculated with *M. oryzae* KEN54-20 (*AVR-Rmo1*) and *avr-Rmo1* mutants M43 and M61. *M. oryzae* isolate KEN54-20 is avirulent on Baronesse, whereas *avr-Rmo1* mutants M43 and M61 are virulent. Ectopic integration of *PWL2* driven by native promoter (*pPWL2:PWL2*) complements the phenotype of mutants M43 and M61. Susceptible control, Nigrate. **(B)** *Mla3* recognizes *PWL2* in transgenic Golden Promise + *Mla3.* Transgenic Golden Promise + *Mla3* (T_1_-4 T_2_) spot inoculated with *M. oryzae* isolate KEN54-20 (*AVR-Rmo1*), *avr-Rmo1* mutants M43 and M61, and *avr-Rmo1* mutants M43 and M61 transformed with *PWL2* driven by its native promoter (*pPWL2:PWL2*). Susceptible control, Golden Promise. **(C)** Weeping lovegrass spray infected with *M. oryzae* isolates with and without *AVR-Rmo1*. **(D)** Baronesse (*Mla3*) spot inoculated with 17 *M. oryzae* isolates representing *PWL2* natural variation. The isolate Ina87T-56A carries both *PWL2* and *pwl2-2*. Susceptible control, Nigrate. Phenotypes are representative of three biological replicates with three technical replicates.

### Conserved recognition specificity of *PWL2* in barley and weeping lovegrass

*PWL2* conditions the ability of rice-infecting *M. oryzae* isolates to infect weeping lovegrass (*Eragrostis curvula*) and is a major determinant of host-species specificity (Kang *et al*., 1995; Sweigard *et al*., 1995). To confirm loss of *PWL2* function, we tested whether KEN54-20 *Δpwl2* mutants were virulent on weeping lovegrass due to loss of *PWL2*. Weeping lovegrass was resistant to spray inoculation of wild-type KEN54-20, yet infection of two independent mutant lines (M43 and M61) resulted in susceptible lesions and restriction of plant growth (**Figure 7c**). Ectopic transformation of *PWL2* in the independent mutants (M43*+pPWL2:PWL2* and M61*+pPWL2:PWL2*) made them avirulent on *E. curvula* (**Figure 7c**). *PWL2* is part of the *PWL* multi-gene family and is highly prevalent across isolates (Kang *et al*., 1995; Sweigard *et al*., 1995; Valent *et al*., 1986). *pwl2-2* encodes an allele of *PWL2* that contains a single non-synonymous (D90N) substitution resulting in the loss of recognition and subsequent virulence on weeping lovegrass (Kang *et al*., 1995; Schneider *et al*., 2010; Sweigard *et al*., 1995). To test whether the same non-synonymous mutation would abolish recognition by *Mla3*, diverse *M. oryzae* isolates were inoculated on Baronesse (**Figure 7d**). Seven *M. oryzae* isolates carrying wild-type *PWL2* were avirulent on Baronesse (*Mla3*) (**Figure 7d**). The *M. oryzae* isolate Ina168, known to lack *PWL2*, was virulent on Baronesse. In agreement with the previous reports on weeping lovegrass (Sweigard *et al*., 1995), *M. oryzae* isolates carrying the *pwl2-2* allele (PWL2 D90N) were not recognized by *Mla3* and therefore virulent on Baronesse. The isolate Ina87T-156A, which carries both *PWL2* and *pwl2-2,* was avirulent on Baronesse due to recognition of the wild type *PWL2.* When considered together, these results provide evidence that recognition of *PWL2* is conserved in barley and weeping lovegrass, and specificity of recognition is maintained in both grass species.

### Weeping lovegrass lacks an *Mla* ortholog

The resistance gene in weeping lovegrass that recognizes *PWL2* is unknown. To determine whether an *Mla* ortholog is present in weeping lovegrass and related grass species, we constructed a maximum likelihood phylogenetic tree using protein sequence for nucleotide binding (NB) domains from NLRs of seven grass species: barley (*H. vulgare*) (Mascher *et al*., 2021), weeping lovegrass (*E. curvula*) (Carballo *et al*., 2019), *Brachypodium distachyon* (Vogel *et al*., 2010), rice (*Oryza sativa*) (Goff *et al*., 2002), maize (*Zea mays*) (Schnable *et al*., 2009), *Setaria italica* (Bennetzen *et al*., 2012), and *Sorghum bicolor* (Paterson *et al*., 2009). These species have high quality genomes and annotations and are balanced in representation between PACMAD and BOP clades. Superposition of previous clade classifications found that the *RGH1* (*Mla*) gene family is in the C17 clade (**Supplemental Figure 9**) (Bailey *et al*., 2018). Phylogenetic analysis of the C17 clade found 34 NLRs from *B. distachyon*, 36 NLRs from barley, 36 NLRs from rice, 30 NLRs from *S. bicolor*, 49 NLRs from *S. italica*, 25 NLRs from *Z. mays*, and 101 NLRs from weeping lovegrass in the C17 clade. While bootstrap support exists for many subclades that found orthologous NLRs from the majority of grasses, the subclade including RGH1 only had members from barley and *B. distachyon* (**Supplemental Figure 10ab**). Thus, the RGH1 family and related NLRs are predominantly absent in the evaluated sequenced species from the PACMAD clade, in which weeping lovegrass resides. Therefore, recognition of *PWL2* in *E. curvula* is likely conferred by a resistance gene outside of the *RGH1* family.

## Discussion

The majority of NLRs confer resistance in a pathogen species- or isolate-specific manner. *Mla* alleles have been well studied for isolate-specific resistance to *Bgh* through the functional divergence of alleles (Jørgensen & Wolfe, 1994; Saur *et al*., 2019; Seeholzer *et al*., 2010). However, while resistance to multiple pathogens has been mapped to the *Mla* locus across barley haplotypes, many of the underlying causal genes have yet to be identified due to the limitations of suppressed recombination across the locus and the inability to assemble the region with current genomic tools. Homologs of *Mla*, *Sr33* in wheat (*Triticum aestivum*) and *Sr50* in rye (*Secale cereale*), confer disease resistance to diverse races of stem rust pathogen *Puccinia graminis* f. sp. *tritici*, including the virulent isolate TTKSK (Chen *et al*., 2017; Periyannan *et al*., 2013). *Mla*, *Sr33*, and *Sr50* highlight the potential for orthologous genes to evolve new and different pathogen specificities, expanding the breadth of recognition of the *Mla* gene family across a diversity of ascomycete and basidiomycete pathogens. In comparison, *Mla8* (=*Rps7*) was shown to confer resistance to both *Bgh* and *Puccinia striiformis* f. sp. *tritici* (Bettgenhaeuser *et al*., 2021). *Rps7* recognition is specific to the non-adapted pathogen *P. striiformis* f. sp. *tritici* and is ineffective against the adapted barley stripe rust *P. striiformis* f. sp. *hordei* (Bettgenhaeuser *et al*., 2021). Here, we confirm that the barley NLR Mla3 recognizes the effector Pwl2 from *M. oryzae* and show that this single NLR is capable of recognition of two taxonomically diverse pathogens. Mla3 specifically recognizes the host-specificity determinant Pwl2, as loss of the *PWL2*-encoded effector permits successful infection by *M. oryzae*. Mutation of the aspartic acid residue in position 90 of Pwl2 is sufficient to abolish recognition, as shown by the allele *pwl2-2* (Pwl2 D90N), demonstrating the specificity of NLR-effector interaction. Therefore, *Mla3* has evolved the capacity to recognize host-adapted effectors from *Bgh* and convergently recognizes an important host-determining effector from *M. oryzae*, further expanding the multiple pathogen recognition of the *Mla* locus.

As a species, *M. oryzae* is known to infect a wide range of cultivated and wild grass plant species; however, individual isolates have a narrow host range due to incompatibility factors present in the plant (Couch & Kohn, 2002; Gladieux *et al*., 2018; Jacob, 2021; Ou, 1985). The effector gene *AVR1*-*CO39* was lost during the early evolution of the rice-specific subgroup of *M. oryzae*, allowing it to infect rice cultivars in which the corresponding *R* gene *Pi*-*CO39(t)* is present (Tosa *et al*., 2005). Recently, the emergence of the wheat infecting lineage of the blast fungus, the *M. oryzae Triticum* pathotype, was found to be due to the sequential loss of specific effectors encoded by *PWT3* and *PWT4* (Inoue *et al*., 2017). The corresponding wheat *R* genes *Rwt3* and *Rwt4* were responsible for incompatibility of *M. oryzae* during infection and their separation facilitated *M. oryzae* step-wise adaptation to become a pathogen of wheat (Inoue *et al*., 2017). Members of the *PWL* (*Pathogenicity toward Weeping Lovegrass*) multi-gene family—*PWL1, PWL2, PWL3*, and *PWL4*—contribute to host specialization of *M. oryzae* pathotypes (Kang *et al*., 1995; Sweigard *et al*., 1995; Valent *et al*., 1986). *PWL1*, first identified from finger millet (*Eleusine coracana*) isolates, and *PWL2*, first identified in rice isolates, prevent infection on weeping lovegrass (*Eragrostis curvula*) and therefore condition host-specificity (Kang *et al*., 1995; Sweigard *et al*., 1995). Small amino acid changes in the Pwl2 effector are sufficient to abolish recognition and allow virulence (Schneider *et al*., 2010), yet *PWL2* displays low genetic variation within populations across geographic locations and the majority of isolates pathogenic on rice contain *PWL2* (Huang *et al*., 2014; Sirisathaworn *et al*., 2017; Sweigard *et al*., 1995). Widespread prevalence of *PWL2* in rice-infecting lineages would be facilitated by lack of resistance genes to *PWL2* in rice. A single amino acid change of D90N for example—found in the *pwl2-2* allele—is sufficient to abolish recognition (Schneider *et al*., 2010; Sweigard *et al*., 1995). As *PWL2* is a mobile effector (i.e. can move between cells) (Khang *et al*., 2010; Sirisathaworn *et al*., 2017), sequence divergence may be constrained by functional requirements. Recognition of *PWL2* may occur through this required structure, or via detection of effector-induced changes to the host environment that are vital for virulence. As yet, the resistance gene in weeping lovegrass that recognizes *PWL2* has not been identified and the mechanism underlying recognition remains unknown.

In a plant species, broad recognition specificity can be maintained via allelic series of NLRs within populations where individual alleles recognize different effectors (Brown & Tellier, 2011; Dangl & Jones, 2001). The allelic flax rust (*Melampsora lini*) resistance gene *L* was the first to be described (Flor, 1956) and the additional *K, M, N*, and *P* loci encode further closely linked or allelic genes (Ravensdale *et al*., 2011). The *A. thaliana RPP8* gene family encodes NLRs with diverse resistance specificities, with alleles conferring resistance to the oomycete *H. arabidopsidis* (*RPP8*); *Turnip crinkle virus* (*HRT*), and *Cucumber mosaic virus* (*RCY1*) (Cooley *et al*., 2000; Kuang *et al*., 2008; McDowell *et al*., 1998; Takahashi *et al*., 2002). It is hypothesized that unequal crossovers that generate chimaeras between alleles are a driving force for new *RPP8* specificities (Ding *et al*., 2007; McDowell *et al*., 1998). Similar to the *Mla* alleles of barley and *Bgh*, the wheat *Pm3* alleles recognize *B. graminis* f. sp. *tritici* (*Bgt;* wheat powdery mildew) *AvrPm3* effectors which are sequence unrelated but structurally similar (Bourras *et al*., 2019; Bourras *et al*., 2018). Direct NLR-effector interaction can drive co-evolution and diversification of NLR and effector repertoires, through opposing selection on NLRs to maintain effector recognition and selection on effectors to avoid triggering resistance (Saur *et al*., 2021; Van der Hoorn *et al*., 2002). However, evasion can be mitigated by NLRs that recognize conserved effectors or protein structures. For example, many bacterial pathogen species contain the effector AvrRpt2 which is important for virulence. This observation, alongside the conserved recognition of AvrRpt2 across diverse plant species, supports the hypothesis that AvrRpt2 or AvrRpt2-like effectors have been present in pathogens throughout evolutionary history (Mazo-Molina *et al*., 2020). There may be a limitation on the functional and structural variation available to effectors and processes that are vital for pathogenesis.

Reciprocal domain swaps between *Mla1* and *Mla6* revealed that the LRR domain was sufficient for recognition of cognate *AVR_a_* (Shen *et al*., 2003) and sequence comparison of alleles confirmed sites of positive selection in the LRR domain (Seeholzer *et al*., 2010). Further interaction studies between Mla10 and Mla22 with AVR_a10_ and AVR_a22_ showed recognition specificity is largely determined by the LRR domain present in Mla, and few residues in the AVR proteins (Bauer *et al*., 2021). Direct recognition has been described for *Bgh* effectors AVR_a1,_ AVR_a7_, AVR_a10_, AVR_a13_, and AVR_a22,_ and the *Mla* alleles, Mla1, Mla7, Mla10, Mla13, and Mla22 respectively (Lu *et al*., 2016; Saur *et al*., 2019). Recognition is highly specific as only the interaction of the *Mla* allele with the matching *Bgh AVR_a_* activates cell death in barley protoplasts and heterologous systems (Chen *et al*., 2017; Saur *et al*., 2019). Known *Bgh AVR_a_* effectors are sequence unrelated, apart from the allelic AVR_a10_, and AVR_a22,_ and are predicted to share a conserved structural fold (Bauer *et al*., 2021; Saur *et al*., 2019). AVR_a6_ belongs to the family of catalytically inactive RNAse-Like Proteins expressed in Haustoria (RALPHs) and similar RNAse-like folds are present in AVR_a1,_ AVR_a7_, AVR_a10_, AVR_a13_, and AVR_a22_. A common structural scaffold shared by the RALPH effector family may therefore be driving diversification of Mla alleles (Bauer *et al*., 2021). In comparison, the *P. graminis* f. sp. *tritici* effector AVR-Sr50 directly binds the Mla homolog Sr50 and its predicted structure does not resemble known *Bgh* AVRs (Chen *et al*., 2017). For *Mla3*, AVR_a3_ has not yet been identified. AlphaFold structure prediction of PWL2 (Jumper *et al*., 2021) and subsequent structural comparisons on the Dali server (Holm, 2022) show that PWL2 does not contain an RNAse-like fold and belongs to the group of MAX effectors (*Magnaporthe oryzae* Avrs and ToxB), which is an expanded family of sequence unrelated but structurally similar effectors in *M*. *oryzae* (de Guillen *et al*., 2015). Using structural predictions of *Bgh* proteins, *Bgh* was not predicted to have effectors clustering within the MAX effector family (Seong & Krasileva, 2022).

An important question arising from this study remains: how do sequence similar *Mla* alleles recognize structurally distinct effectors from diverse pathogen species? The recognized effectors from *Bgh*, *P. graminis* f. sp. *tritici*, and now *M. oryzae* represent different structural families. Recent work has shown that Mla alleles and the Mla homolog Sr50 recognize effectors through direct recognition and is supported through yeast-two-hybrid assay, *Nicotiana benthamiana* infiltration assay, and barley and wheat protoplast systems where co-expression of Mla and *Bgh* AVRs is sufficient to trigger cell death (Bauer *et al*., 2021; Chen *et al*., 2017; Saur *et al*., 2019). It is unclear if the specificity of recognition is determined by shared or distinct regions of the LRR domain across alleles for each effector. Comparing the structure and binding interfaces of *PWL2* and *AVR_a3_*—once identified—will provide an initial step to elucidate the mechanism of recognition. Interaction studies between *Mla3* and each effector will confirm if additional components are needed for recognition and signaling. If *AVR_a3_* and *PWL2* are directly recognized by *Mla3* due to a shared conformation, this could represent a core structural feature of pathogen effectors conserved across taxonomically diverse species. This may reveal a structural signature to guide effector discovery and NLR engineering for increased recognition and resistance.

## Materials and Methods

### Plant material

Barley accessions were obtained from United States Department of Agriculture Germplasm Resource Information Network (Aberdeen, ID, USA), Oregon State University (Corvallis, OR, USA), John Innes Centre (Norwich, UK), Wageningen University & Research (Wageningen, Netherlands), and CSIRO (Canberra, Australia). All barley accessions were subjected to single seed descent before carrying out subsequent experiments. *E. curvula* was obtained from Star Seed, Inc. (Kansas, USA).

### Recombination screen

Barley accessions Baronesse and BCD47 were crossed and allowed to self-pollinate to generate a founder F_2_ population. For the Baronesse x BCD47 population, seedlings were germinated in John Innes Peat & Sand Mix (85% Fine Peat, 15% Grit, 2.7 kg/m^3^ Osmocote 3-4 months, Wetting Agent, 4 kg/m^3^ Maglime, 1 kg PG Mix). Leaves were sampled at second leaf emergence, DNA extracted, and individuals genotyped for recombination events. Recombinants were transferred to larger pots in John Innes Cereal Mix (40% Medium Grade Peat, 40% Sterilised Soil, 20% Horticultural Grit, 1.3 kg/m^3^ PG Mix 14-16-18 + Te Base Fertiliser, 1 kg/m^3^ Osmocote Mini 16-8-11 2 mg + Te 0.02% B, Wetting Agent, 3 kg/m^3^ Maglime, 300 g/m^3^ Exemptor) and grown in a greenhouse under natural conditions.

Recombination events in a 22.9 cM region on chromosome 1H including *Mla3* were identified in the Baronesse x BCD47 F_2_ population using a total of 1,152 individuals (2,304 gametes). Following single-seed descent, DNA was extracted from leaf tissue of F_2_ and F_2:3_ recombinants using a CTAB-based protocol (Stewart & Via, 1993). Briefly, 3 g of leaf tissue were ground on liquid N_2_ and homogenized with 20 mL of CTAB extraction buffer (2% CTAB, 100 mM Tris-HCl pH8.0, 20 mM EDTA pH8.0, 1.4 M NaCl, 1% β-Mercaptoethanol). Samples were incubated for 30 min at 65°C followed by two chloroform extractions and ethanol precipitation. DNA was then resuspended in 1x TE, 50 μg/mL Rnase A solution and incubated for 1 h at 37°C. DNA was subsequently precipitated with 2.5 volumes of ice-cold 95% ethanol and resuspended in 1x TE. Quantification of DNA samples was performed using a Nanodrop spectrophotometer (Thermo Scientific) and the Qubit dsDNA HS Assay Kit (Molecular Probes, Life Technologies).

Genetic markers designed for the barley oligonucleotide pool assay (BOPA1) panel were converted to Kompetitive allele specific PCR (KASP) markers, which are also SNP based (**Supplemental Data 3**) (Close *et al*., 2009). Briefly, KASP SNP genotyping uses two competitive, allele-specific forward primers and one common reverse primer for allele-specific oligo extension, amplification, and fluorescence output. Genotyping was performed by the John Innes Cetnre Genotyping Facility (Norwich, UK). Genetic maps were created using JoinMap v4 was used using default parameters (Van Ooijen, 2006). Genetic distances were estimated using the Kosambi mapping function. Integrity of the genetic map was evaluated through comparison with the current OPA consensus genetic map of barley (Muñoz-Amatriaín *et al*., 2011) and using Rstudio (Version 1.1.463) and R/qtl package (Version 1.44.9) (Broman *et al*., 2003).

### *RNAseq and* de novo *transcriptome assembly*

First and second leaf tissue was harvested at 10 days after sowing of barley accessions grown in the greenhouse. Tissue was flash frozen in liquid nitrogen and stored at −80 °C. Tissue were homogenized into a fine powder in liquid nitrogen-chilled pestle and mortars. RNA was extracted, purified, and quality assessed as described by (Dawson *et al*., 2016). RNA libraries were constructed using Illumina TruSeq RNA library preparation (Illumina; RS-122-2001). Barcoded libraries were sequenced using either 100 or 150 bp paired-end reads. Library preparation and sequencing was performed by Earlham Institute (Norwich, United Kingdom), Novogene (Cambridge, UK), and BGI (Shenzhen, China). Quality of all RNAseq data was assessed using FastQC (0.11.5) (Andrews, 2010). Trinity (2.4.0) (Grabherr *et al*., 2011) was used to assemble *de novo* transcriptomes using default parameters and Trimmomatic (Bolger *et al*., 2014) for read trimming. Genes of interest were identified in assemblies using BLAST+ (v2.2.9) (Camacho *et al*., 2009).

### Long-range assembly of Baronesse chromosome 1H

Long-range sequencing and assembly were carried out as described by (Holden *et al*., 2022). Briefly, chromosome flow sorting of Baronesse chromosome 1H and preparation of its DNA was performed using the methods described by Doležel *et al*. (2021) and Zwyrtková *et al.* (2021) with an estimated purity of 82.8 to 92.3% in the sorted fractions (**Supplemental Figure 11**). Chromosomal high molecular weight (HMW) DNA, Chicago libraries and sequencing (Dovetail Genomics, Santa Cruz, CA, USA), assembly in Meraculous (v2.0.3), and final scaffolding in HiRise were performed as described in (Thind *et al*., 2017). Shotgun sequencing was carried out using two insert sizes, estimated at 221 bp and 454 bp, with 266.3 and 246.9 million paired end reads, respectively. An initial assembly using Meraculous had length 450.7 Mb on 40,855 scaffolds. The HiRise assembly had 454.5 Mb on 2,009 scaffolds.

### Sequence capture and PacBio SMRT sequencing

Sequence capture and PacBio SMRT sequencing of NLR encoding genes (RenSeq) were carried out according to (Witek *et al*., 2016). A custom Daicel Arbor Biosciences Mybaits bait library was previously designed on the barley NLR gene space including the entire Mla locus from barley accession Morex (TSLMMHV1) (Brabham *et al*., 2018). PacBio circular consensus sequencing was performed at the Earlham Institute (Norwich, UK). *De novo* assembly was performed using Geneious (v10.2.3) using custom sensitivity parameters for assembly: don’t merge variants with coverage over approximately 6, merge homopolymer variants, allow gaps up to a maximum of 15% gaps per read, word length of 14, minimum overlap of 250 bp, ignore words repeated more than 200 times, 5% maximum mismatches per read, maximum gap size of 2, minimum overlap identity of 90%, index word length 12, reanalyze threshold of 8, and maximum ambiguity of 4.

### Copy number variation analysis

Copy number analysis was performed using the *k*-mer analysis toolkit (KAT; v.2.4.1) (Mapleson *et al*., 2017). The sequence coverage estimator tool (sect) was used to determine coverage for genomic contigs encompassing *Bpm*, *Mla3*, *RGH2*, and *RGH3*. Default parameters were used including *k*-mer length of 27 bp. *k*-medoids clustering was performed with R (4.1.2) using *pam* from the *cluster* package (2.1.3). Clustering was performed on coverage values between 0 to 1,000.

### Construct development

PCR reactions were performed using GoTaq G2 Flexi DNA polymerase (Promega; Catalogue number M7805), Phusion High Fidelity DNA polymerase (New England Biolabs Ltd; Catalogue number M0530S) and GoTaq Long PCR Master Mix (Promega; Catalogue number M4020). Reaction mixes were set up according to manufactures instructions and performed in a thermal cycler. cDNA was used as the template for cloning of the *RGH1* candidate genes, and gDNA the template for the *RGH2* and *RGH3* construct. Annealing temperatures and elongation times were optimized for reaction based on primer combination and ranged between 52°C – 64°C. PCR reactions were assayed on a 1% agarose gel in TBE or TAE buffer. Gel extraction of fragments was performed with the QIAquick gel extraction kit (Qiagen; Cat No.: 28704) according to the manufacturer’s instructions. Excised and cleaned fragments were A-tailed via incubation at 72°C for 20 mins using GoTaq polymerase and dATPs, cloned via the TOPO XL PCR Cloning Kit (Invitrogen; K7030-20) according to manufacturer’s instructions, and transformed into DH5α *E. coli* competent cells (1 to 2 μL reaction into 50 μL cells). Transformations were placed on ice for 30 mins, heat-shocked at 42 °C for 90 sec, placed on ice for 2 min, recovered in 500 μL L media via shaking at 37 °C for 1 hour; and plated on L media plates in varying dilutions with appropriate selection for overnight growth at 37 °C. Positive clones were identified by PCR using gene specific primers. Plasmids were extracted from positive colonies using 5 mL liquid cultures with the NucleoSpin Plasmid Purification kit (Macherey-Nagel; Ref.: 11932392) and sequenced (GATC; 80 to 100 ng/ plasmid DNA, 5 μM primer). Plasmids were also confirmed through digestion with the restriction enzyme EcoRI (New England Biolabs; Ref.: R3101S) according to the manufacturer’s instructions. Presence of *Mla3Δ6* and differentiation between *Mla3* was assessed via digestion with *Bsp*LI (Thermo Scientific; Cat No.: ER1151) which cuts on the 6 bp indel.

Plant transformation constructs were assembled via multiple PCR fragments into the pBract202 vector backbone. pBract202 was generated by the Crop Transformation (BRACT) team at the John Innes Centre (Norwich, UK) (Smedley & Harwood, 2015). The backbone contains the *npt1* kanamycin resistance gene for bacterial selection and the left border contains the 35S hygromycin selectable marker for plant transformation. Primers for Gibson Assembly consisted of 20 bp fusion from both fragments to be assembled (40 bp total) and were assessed for GC content (~50%) and secondary structures using mfold (Zuker, 2003). Constructs were assembled via using Gibson Assembly (Gibson *et al*., 2009) with a Gibson Assembly master mix (New England Biolabs; Ref.: E2611). Briefly, multiple overlapping gene fragments are designed and amplified via PCR; appropriate overlaps are unique ~18 bp overhangs. Overlaps were added using the high fidelity Phusion polymerase. PCR products were digested with *Dpn*I (New England Biolabs; Ref.: R0176S) to remove circular DNA. Fragments were resolved with gel electrophoresis (1% agarose in 1x TAE buffer) and excised using the Zymoclean Gel DNA Recovery Kit (Zymo Research; Ref.: D4008) for elution of high-concentration ultra-pure DNA. Fragments were added to the reaction tube in appropriate dilutions and ratios according to molecular weight, as outlined in the manufacturer’s instructions. The reaction tube was incubated at 50°C for 1 h – fragments trimmed by an exonuclease creating single-stranded 3’ overhangs that anneal with their complementary counterparts, DNA polymerase extends 3’ ends of annealed fragments, and sealed with a DNA ligase. Competent *E. coli* DH5α cells were transformed with the successful construct as outlined above. Construct visualization, primer development, and assessment was performed using the software Geneious (Version 9.0.5). Construct maps are shown in **Supplemental Figure 6**.

### Plant transformation

Constructs were transformed into *Agrobacterium tumefaciens* AGL1 containing pSoup via electroporation (~100 ng plasmid into 50 μl cells), recovered in 500 μL L medium via shaking at 28°C for 2 h, and grown on L media plates with appropriate selection for three days. Barley accession Golden Promise was transformed using *Agrobacterium*-mediated transformation based on the approach described by (Hensel *et al*., 2009). Assessment of insert copy number of transgenic lines was performed by iDna Genetics Ltd (Norwich, UK) using real-time PCR assaying the presence of the hygromycin resistance gene.

### B. graminis *propagation and inoculation*

*Bgh* isolate CC148 was obtained from James Brown (John Innes Centre, Norwich, UK). Seedlings for phenotypic assays were germinated in John Innes Cereal Mix at 18°C under a 16-h light and dark cycle. Seedlings were transferred to a containment greenhouse set at 18°C day for 16 h and 12°C night for 8 h with supplemental lighting from 400W HQI (metal halide lamps). *Bgh* inoculations were carried out on seedlings at emergence of the second leaf. *Bgh* inoculum was maintained on the susceptible barley accession Manchuria. Inoculation was carried out by laying pots on their side and gently shaking infected leaves over both sides. Phenotyping was carried out as described in (Bettgenhaeuser *et al*., 2021).

### M. oryzae *propagation and inoculation*

Protocols for culturing and inoculation of *M. oryzae* were similar as described by (Jia *et al*., 2003; Parker *et al*., 2008). *M. oryzae* isolates were maintained on Potato Dextrose Agar medium at 24°C and as frozen stocks of dried mycelium on Whatman filter paper (GE Healthcare Whatman™ Qualitative Filter Paper: Grade 1 Circles, Fisher Scientific UK) at - 20°C. Hyphal tips were transferred to oatmeal agar (20 g oatmeal, 10 g agar, 2.5 g sucrose, addition of ddH20 to 500 ml) plates (deep petri dish 100 x 20 mm) for the production of spores (conidia) and incubated for 10-15 days at 24°C. To increase spore production, plates were used for a second time after washing and a further 10-15 day incubation. For *M. oryzae* inoculation, seedlings were germinated in John Innes Cereal Mix (40% Medium Grade Peat, 40% Sterilised Soil, 20% Horticultural Grit, 1.3 kg/m^3^ PG Mix 14-16-18 + Te Base Fertiliser, 1 kg/m^3^ Osmocote Mini 16-8-11 2 mg + Te 0.02% B, Wetting Agent, 3 kg/m^3^ Maglime, 300 g/m^3^ Exemptor). Seedlings were germinated and grown in a controlled environment at 25°C under a 16-h light and dark cycle. *M. oryzae* conidia were collected by the addition of 8 mL dH_2_O to the oatmeal agar plates and gentle scraping with the tip of a 1.5 mL microcentrifuge tube. Suspension was poured and filtered through Miracloth (Merck Chemicals, Ref.: 475855-1r) and collected in a 50 mL Corning tube. Spore concentration was counted via haemocytometer and adjusted to 1 x 10^5^ spores per mL in 0.1% gelatin or dH_2_O with 0.01% Tween 20 (Merck Chemicals, CAS Number: 9005-64-5).

*M. oryzae* spot inoculations were carried out on detached leaves on agar (2.5 g Agar-agar (Fisher, CAS 9002-18-0); 50 mL benzimidazole (1 g / 1 L H_2_O stock solution); 450 mL H_2_O). Barley was germinated at 25°C under a 16-h light and dark cycle and 1-week old seedlings were used for inoculation at emergence of second leaf. The first leaf was cut and placed on agar inside the boxes. Each leaf was inoculated with 3 to 4 drops of 5 μL of conidal suspension. Boxes were placed at 25°C in a Sanyo growth cabinet and maintained under continuous light for the first 24 h. After 24 h droplets were removed from the leaves using sterile Miracloth and boxes returned to 25°C in a 16-h light and dark cycle. Detached leaves were monitored for development of lesions and phenotypes were recorded 7 days post inoculation (dpi). Phenotypes were scored as resistant on a scale of 0 = complete resistance; 1 = small brown resistant spots; 2 = susceptible larger eyespot lesions; 3 = larger spreading lesions; 4 = hyper susceptibility.

*M. oryzae* spray inoculations were carried out on whole 1-week old seedlings at emergence of the second leaf. Barley was germinated 25°C under a 16-h light and dark cycle with 9 seeds placed in a 9 cm^2^ pot, with 8 pots in a tray. Each tray was sprayed with ~5 mL of conidial suspension using a 20 mm atomizer spray bottle. Trays were placed in polythene autoclave bags tied with tape and placed inside a Sanyo cabinet at 25°C under a 16-h light and dark cycle. Bags remained covering the plants until phenotyping due to containment requirements. Disease symptoms were recorded 7 dpi and first leaves scored on a similar phenotypic scale to spot inoculations. A similar protocol was followed for spray inoculations of 10-day old weeping lovegrass (*Eragrostis curvula*) plants.

### M. oryzae *mutagenesis*

Mutagenesis of *M. oryzae* isolate KEN54-20 was performed on the conidial suspension using ultraviolet light (UV). Spore concentration was adjusted to 1 x 10^5^ spores per mL and placed inside a petri dish until the solution was just covering the entire base of the dish – a shallow depth is required to ensure even UV light penetration. The open petri dish was placed inside a UV Stratalinker 2400 (Stratagene) and exposed to set UV light. A dosage curve was generated to assess the UV dose at which spore death was at 50%, for KEN54-20 this was 20 sec. The UV light exposed conidial suspension was used for spray-based inoculation. Wild-type Baronesse was used for the isolation of KEN54-20 gain-of-virulence mutants. Lesions were isolated from leaves 7 dpi, sterilized in ethanol for 30 sec, and placed on Potato Dextrose Agar (10g PDA, 6.25g agar per liter of water). *M*. *oryzae* growth was sampled from each lesion, cultured, and re-inoculated onto Baronesse to confirm gain-of-virulence.

### M. oryzae *transformation*

Transformation of *M. oryzae avr-Rmo1* isolates was performed as previously described (Talbot *et al*., 1993). Briefly, the region of fungal active growth was cut from the surface of a CM agar plate, blended in 50 mL of CM liquid media, and incubated for 48 h at 25°C and 120 rpm. The culture was filtrated through two layers of sterile Miracloth™ and mycelia was gently resuspended and digested with Glucanex™ in 0.7 M NaCl (pH 5.5, filter sterilized). Protoplasts were generated after incubation for 3 h at 25°C and gentle shaking at 75 rpm. The digested mycelium was filtered through sterile Miracloth™ and protoplasts were collected and centrifuged at 3500*xg* and 4°C for 10 min. Protoplasts were washed with STC buffer (1.2 M sorbitol, 10 mM Tris-HCl pH 7.5, 10 mM CaCl_2_), centrifuged at 3500*xg* for 10 min and resuspended in 150 μL of STC buffer. Transformation was carried out by mixing the protoplasts with 4 μg of the vector pCB1532∷*pPWL2:PWL2:tPWL2*, incubating at room temperature for 15 min, and subsequently adding 1 mL of PTC buffer (60% PEG 4000, 10 mM Tris-HCl pH 7.5, 10 mM CaCl_2_). The mix was let to stand for 5 min at room temperature, added to BDCM liquid media (0.8 M sucrose, 1.7 g L-1 yeast nitrogen base without amino acids and Ammonium sulphate, 2 g L-1 Ammonium nitrate, 1 g L-1 asparagine, 10 g L-1 glucose) and incubated overnight at 25°C and 120 rpm. The protoplast culture was added to molten BDCM agar and poured onto plates. Selective BDCM media (BDCM media lacking glucose) with 1% agar and sulfonylurea (150 μg/mL Chlorimuron ethyl) was added on top as overlay. Plates were incubated at 25°C for 7-10 days until transformed colonies emerged and started to grow. Individual colonies were transferred to BDCM agar plates with 100 μg/mL sulfonylurea and kept for confirmation by PCR and further assays.

### M. oryzae *high molecular weight genomic DNA extraction*

High molecular weight genomic DNA extraction from protoplasts of *M. oryzae* was performed for Oxford Nanopore DNA sequencing of the isolate KEN54-20. Protoplasts were obtained as previously described for *M. oryzae* transformation. Extraction of high molecular weight DNA was carried out as previously described by (Schwessinger & Rathjen, 2017) with some modifications. Briefly, lysis buffer was prepared as follows: 2.5 volumes of autoclaved buffer A (0.35M Sorbitol, 0.1M Tris-HCl, 5mM EDTA pH 8.0), 2.5 volumes of autoclaved buffer B (0.2M Tris-HCL, 50mM EDTA pH8.0, 2M NaCl, 2% CTAB), 1 volume of filter-sterilized buffer C (5% Sarkosyl N-lauroylsarcosine sodium salt), 1 volume of 10% Polyvinylpyrrolidone 40, 1 volume of 10% Polyvinylpyrrolidone 10 and 10 uL of RNAse A (Thermo Fisher). Protoplasts were pelleted and thoroughly resuspended in preheated lysis buffer and incubated under constant rotation for 30 min at room temperature. Proteinase K (New England BioLabs) was added to the sample and further incubated under permanent rotation for 30 min, followed by five min on ice. The sample was mixed with 0.2 volumes of 5M potassium acetate, incubated on ice for no longer than five min and centrifuged at 4°C and 5000x*g* for 12 min. The supernatant was transferred and mixed with one volume of phenol:chloroform:isoamyl alcohol (P:C:I) (25:24:1). After mixing by inversion for two min, the sample was centrifuged at 4°C and 4000x*g* for 10 min. The supernatant was recovered and mixed once more with one volume of P:C:I, followed by centrifugation at 4°C and 4000x*g* for 10 min to separate the organic phase and remove proteins. The supernatant was mixed by inversion with 0.1 volumes of 3M sodium acetate and then one volume of isopropanol was added. The sample was incubated at room temperature for five min and then centrifuged at 4°C and 8000x*g* for 30 min. The supernatant was discarded and the DNA, visible as a translucent pellet at the bottom, was washed with 70% ethanol and then centrifuged at 5000x*g* for five min. Four more additional washing steps with 70% ethanol were performed, with the final two spins at 13000x*g*. The ethanol was discarded and the DNA pellet was let to air-dry for five min. The DNA was resuspended in molecular grade water and let to dissolve at room temperature. The sample was treated with RNAse A (Thermo Fisher) and column-purified using the Genomic DNA Clean and Concentrator Kit (Zymo Research) according to manufacturer’s instructions. DNA concentration was measured using the Qubit™ dsDNA HS Assay Kit (Thermo Fisher).

### M. oryzae *DNA sequencing*

For genomic DNA Illumina sequencing, mycelia from one-week-old plates of *M. oryzae* isolates KEN54-20 and *avr-Rmo1* mutants were collected, ground with liquid nitrogen to a very fine powder and transferred into 1.5 mL microcentrifuge tube until about two thirds full. 500 uL of CTAB buffer (0.2 M Tris-HCl pH 7.5, 50 mM EDTA, 2 M NaCl, 2% CTAB) pH 7.5 were added and samples were incubated at 65°C for 30 min, shaking every 10 min. Subsequently, 500 uL of chloroform:isoamyl alcohol (24:1) were added and samples were incubated for 30 min under constant shaking at 300 rpm, followed by centrifugation at 16000*xg* for 10 min. The aqueous phase (supernatant) was transferred into a new 1.5 mL microcentrifuge tube and 500 uL of chloroform:isoamyl alcohol (24:1) were added, mixed for five min, and then centrifuged at 16000*xg* for 10 min. The top aqueous phase was transferred to a new tube and 1mL of ice-cold isopropanol was added and mixed. Samples were incubated at −20°C for two h and then centrifuged at 16000*xg* for 10 min. The supernatants were discarded. DNA pellets were allowed to drain for five min and then completely resuspended in 500uL of sterile water. 50 uL of 3 M NaOAc were added with 1 mL of ice-cold 100% ethanol, followed by incubation at −20°C for one hour and centrifugation at 16000*xg* for 20 min. The supernatants were discarded and 400 uL of ice-cold 70% ethanol were added. The samples were centrifuged at 16000*xg* for 5 min, the supernatants were discarded and the pellets were allowed to air-dry. DNA samples were resuspended in 100 uL of TE+RNAse A (Thermo Fisher) and stored at 4°C. Concentration of DNA samples were measured using the Qubit™ dsDNA HS Assay Kit (Thermo Fisher). DNA samples were submitted for library preparation and whole genome sequencing by Illumina to Novogene. The isolate KEN54-20 was sequenced with paired end, 150 bp reads with libraries of 400 bp and 600 bp inserts, and KEN54-20 *avr-Rmo1* mutants were sequenced with paired end, 150 bp reads with libraries of 400 bp inserts.

### Oxford Nanopore DNA sequencing

The gDNA library of *M. oryzae* KEN54-20 was prepared without shearing to maximize sequencing read length. Short DNA fragments were removed with the Short Read Eliminator Kit (Circuloromics®) according to manufacturer’s intructions. DNA repair, end-prep, adapter ligation and clean-up were performed according to the 1D Lambda Control Experiment (SQK-LSK109) protocol provided by ONT. The library was loaded into an R9.4.1 FLO-MIN106 flow cell and MinION sequencing was performed according to ONT guidelines using the ONT MinKNOW software.

### Genome assembly

Base calling of ONT sequencing data was performed with Guppy v3.2.2. Read quality assessment was performed using Pauvre (https://github.com/conchoecia/pauvre) and trimmed using NanoFilt (Li *et al*., 2009). The hybrid assembler MaSuRCA v3.3.3 (Zimin *et al*., 2013) was used to assemble the reference genome of *M. oryzae* isolate KEN54-20 including ONT and Illumina data. Pilon (Walker *et al*., 2014) was used to improve the genome assembly. Alignment of Illumina reads to ONT data was performed using BWA (v0.7.12-r1039; http://bio-bwa.sourceforge.net/). Quality of the assembled and polished genome was assessed using the *k*-mer Analysis Toolkit (KAT; https://github.com/TGAC/KAT). *Ab initio* gene prediction was performed using Augustus (v3.3.2; https://github.com/Gaius-Augustus/Augustus) with the *M. oryzae* species gene model prediction. Genome assembly and annotation completeness were assessed with BUSCO v3 (Waterhouse *et al*., 2018). The *PWL2* region was investigated manually by aligning ONT reads to the reference genome using minimap2 (v2.17-r954-dirty). Illumina reads of *M. oryzae* KEN54-20 wild-type and *avr-Rmo1* mutants were aligned to the genome using BWA. Aligned reads were inspected using IGV (v2.5.3) (Robinson *et al*., 2011). A full set of commands are available on Github (https://github.com/matthewmoscou/AvrRmo1).

### *Identification of common deletions in* M. oryzae avr-Rmo1 *mutants*

The *k*-mer analysis toolkit (KAT; v2.4.1) was used to scan the genome to identify *k*-mers (k=27) that were present in wild-type but absent in all *avr-Rmo1* mutants. A genome scan was performed by counting the number of *k*-mers present in wild-type and absent in all mutants within a window of 10 kb with step size 1 kb.

### Phylogenetic analysis of grass NLRs

To identify NLRs from diverse grass species (**Supplemental Data 4**), InterProScan v5.36-75.0 using default parameters was used to annotate individual protein domains. Proteins annotated with the nucleotide binding domain Pfam family (PF00931) were identified and individual domains extracted from NLRs using the Python script QKdomain_process.py (Bailey *et al*., 2018). Structure-guided multiple sequence alignment of NB domains was performed with MAFFT (v7.481) using DASH (default parameters). NB structures included in the alignment were derived from *Arabidopsis thaliana* ZAR1 (PDB 6J5T) and *Solanum lycopersicum* (tomato) NRC1 (PDB 6S2P). The QKphylogeny_alignment_analysis.py Python script was used to filter the alignment for variable sites represented in at least 40% of proteins and sequences spanning at least 50% of the alignment length (https://github.com/matthewmoscou/QKphylogeny). The phylogenetic tree was constructed using RAxML (v8.2.12) with the JTT amino acid substitution model, gamma model of rate heterogeneity, and 1,000 bootstraps. A convergence test performed using RAxML autoMRE found convergence for both the full NB and C17 clade after 250 bootstraps. iTOL was used for phylogenetic tree visualization and *A. thaliana* ZAR1 was used as outgroups.

## Data Availability

The RNA-seq data used in this study are found in the NCBI database under BioProject codes PRJNA292371, PRJNA376252, PRJNA378334, and PRJNA378723. The genome assembly and sequencing data for barley accession Baronesse chromosome 1H generated in this study have been deposited in the NCBI database under BioProject code PRJNA879438. RenSeq-PacBio of Baronesse raw, circular consensus sequences, and de novo assembly have been deposited in the NCBI database under BioProject code PRJNA422986. Genomic sequencing data and assembly for *M. oryzae* isolate KEN54-20 and *avr-rmo1* have been deposited in the NCBI database under BioProject code PRJNA881958. The sequences of plasmids used for plant transformation in this study have been deposited in the NCBI database with accession codes OP561810 (*Mla3*), OP561809 (*Mla3△6*), and OP561811 (*RGH2*/*RGH3*). All data needed to evaluate the conclusions in the paper are present in the paper and/or the Supplementary Materials. Raw genotypic, phenotypic, and source data for figures and supplemental figures have been deposited on Figshare (https://doi.org/10.6084/m9.figshare.21365532.v1). A material transfer agreement with The Sainsbury Laboratory is required to receive the materials. The use of the materials will be limited to noncommercial research uses only. Please contact M.J.M. regarding biological materials, and requests will be responded to within 60 days.

## Acknowledgements

We greatly appreciate valuable discussions with Sebastian Schornack, Sophien Kamoun, Isabel Saur, Takaki Maekawa, and Paul Schulze-Lefert. Genotyping was supported by Richard Goram (John Innes Centre genotyping facility). Photography was supported by Andrew Davis and Phil Robinson. Assistance in the greenhouse was provided by Sue Banfield and the John Innes Centre Horticultural team. Seed was provided by Rients Niks, Wendy Harwood, Patrick Hayes, Wolfgang Spielmeyer, and the National Small Grains Collection (USDA-ARS). We thank P. Cápal, M. Said, Z. Dubská, J. Weiserová, and E. Jahnová (Institute of Experimental Botany, Olomouc) for assistance with chromosome flow sorting and preparation of chromosome DNA.

---

**Supplemental Figure 1.**
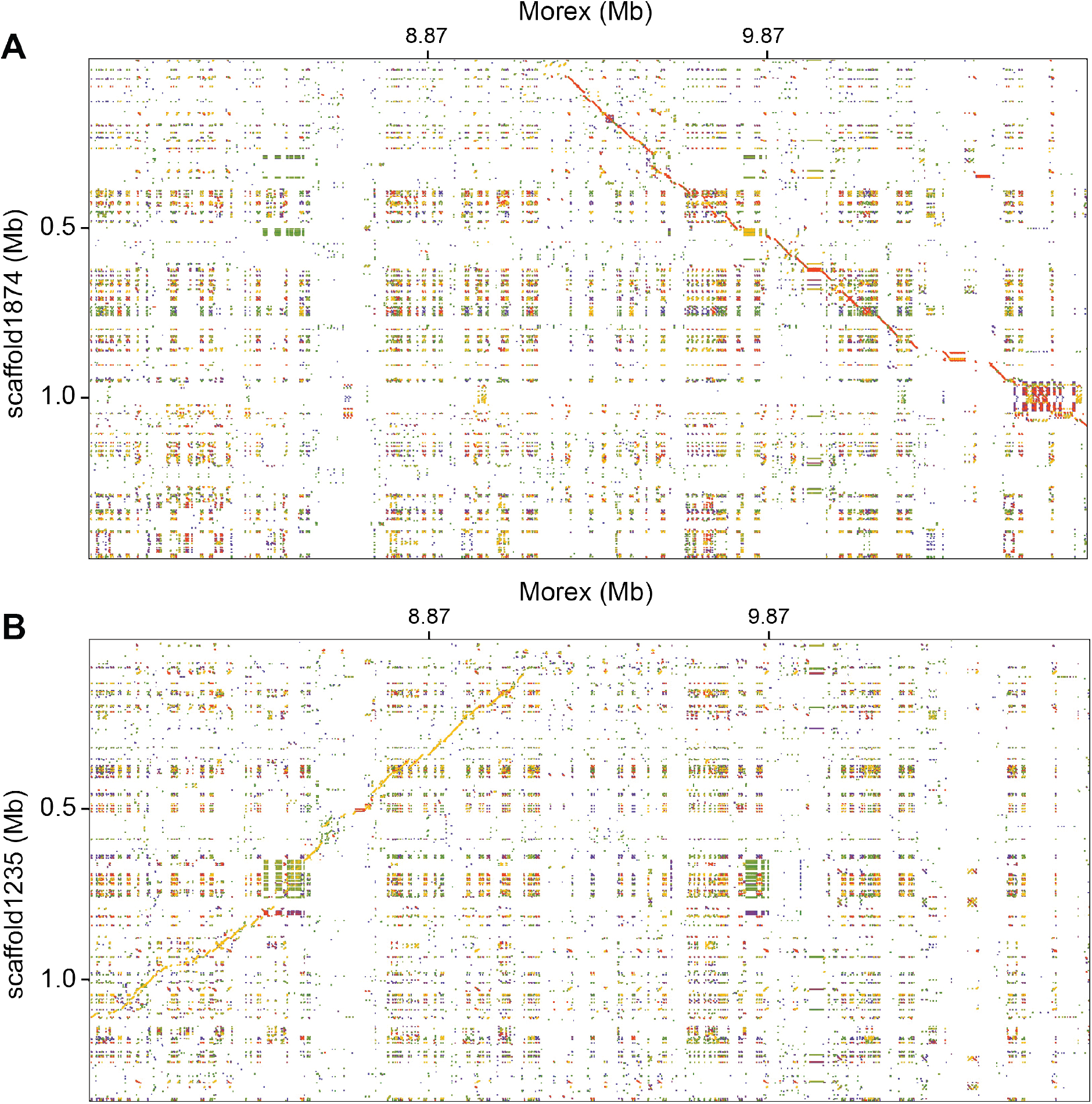
Genomic regions flanking the *Mla* locus are conserved between barley accessions Morex and Baronesse. Dotplot alignments of barley accession Baronesse (A) scaffold1874 and **(B)** scaffold1235 versus the genomic region encompassing *Mla* in barley accession Morex on chromosome 1H (7.87 to 10.81 Mb).

**Supplemental Figure 2.**
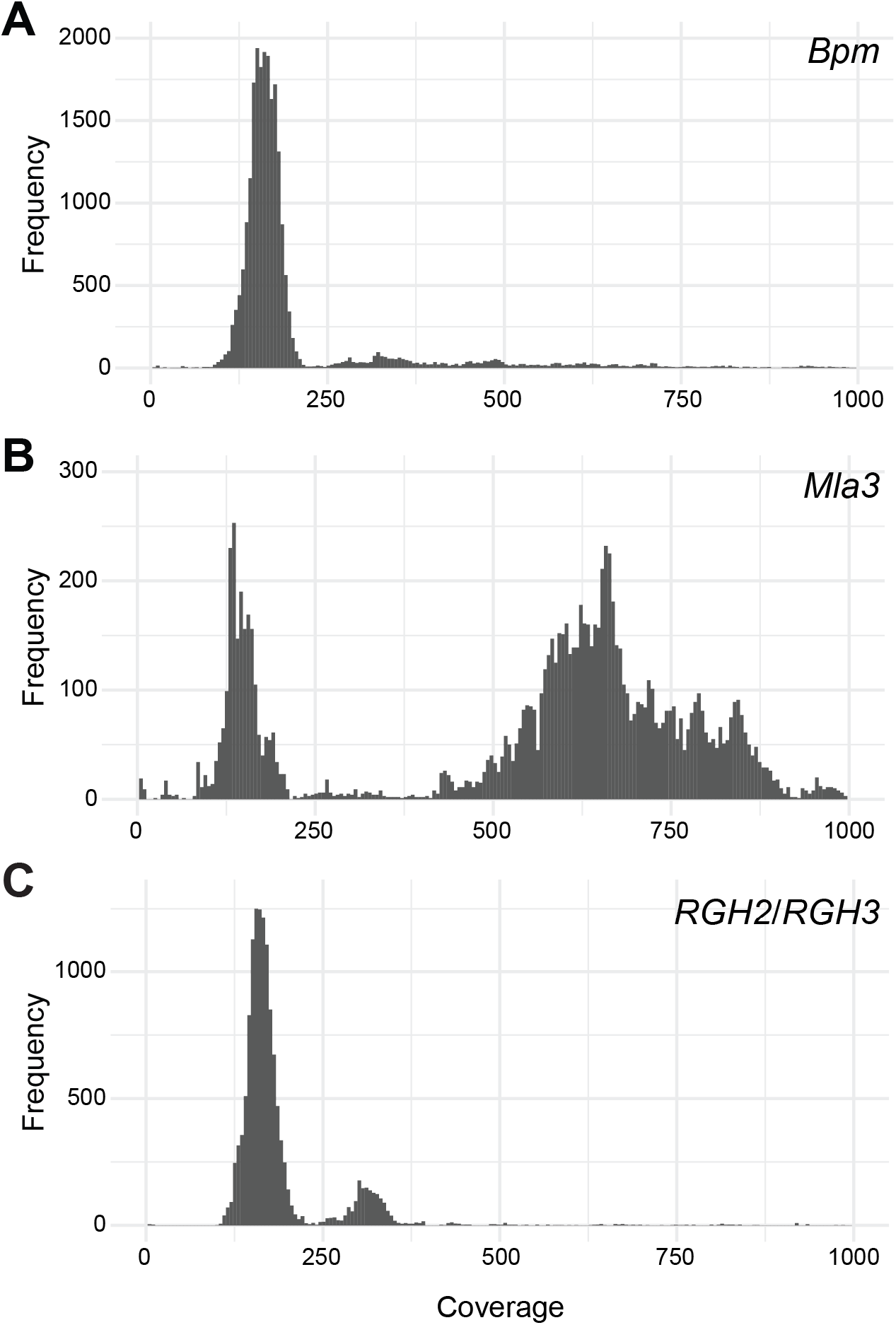
*k*-mer coverage distribution for *Bpm*, *Mla3*, and *RGH2*/*RGH3*. *k*-mer coverage was determined using the K-mer Analysis Toolkit (KAT) for the genomic sequence of (A) *Bpm*, (B) *Mla3*, and (C) *RGH2*/*RGH3*.

**Supplemental Figure 3.**
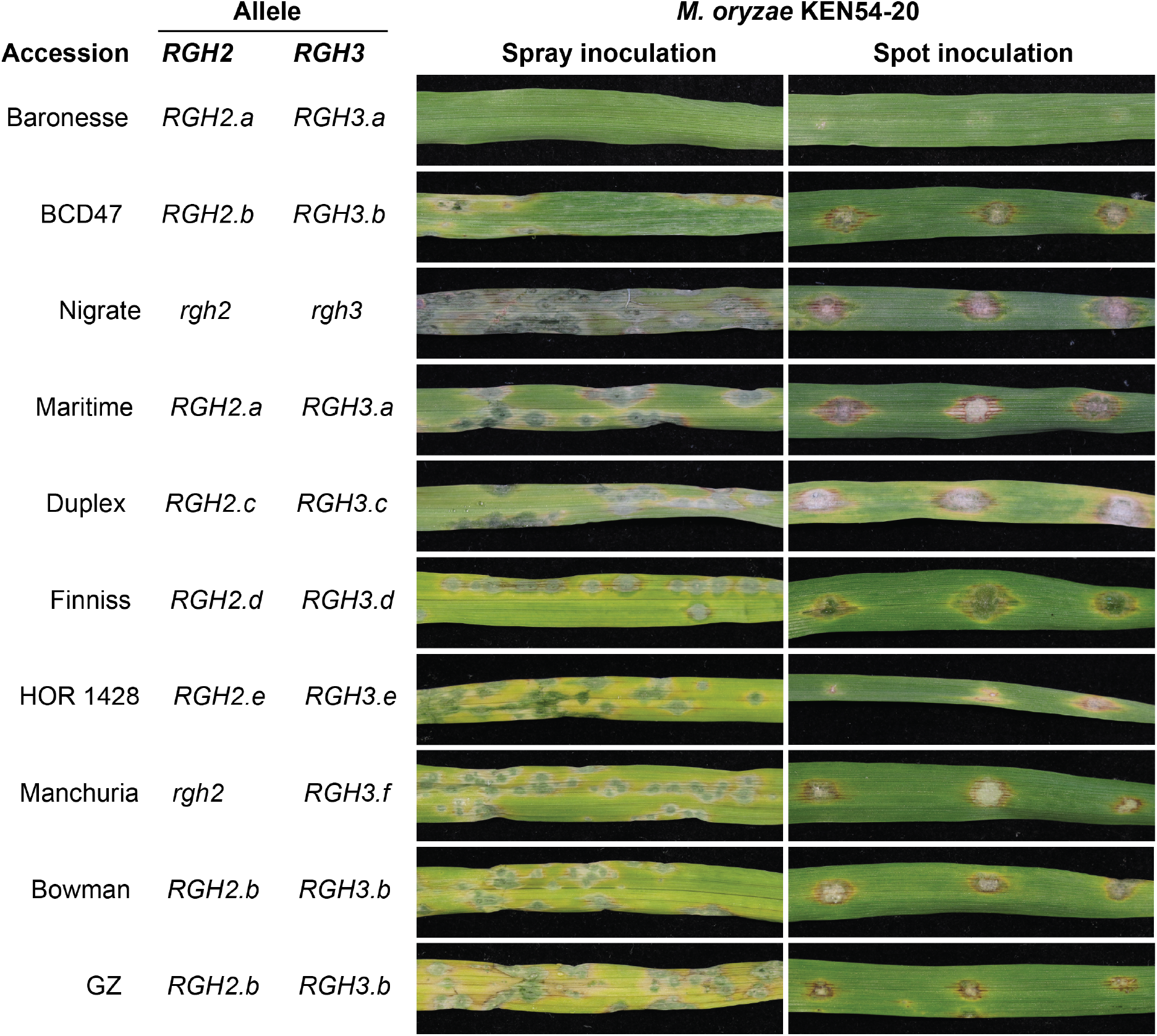
Barley accessions carrying identical *RGH2* and *RGH3* alleles in Baronesse are susceptible to *M. oryzae* KEN54-20. Diversity panel of barley accessions inoculated with *M. oryzae* KEN54-20 using spot and spray-based inoculation protocol. Phenotypes are representative of three biological replicates with four technical replicates.

**Supplemental Figure 4.**
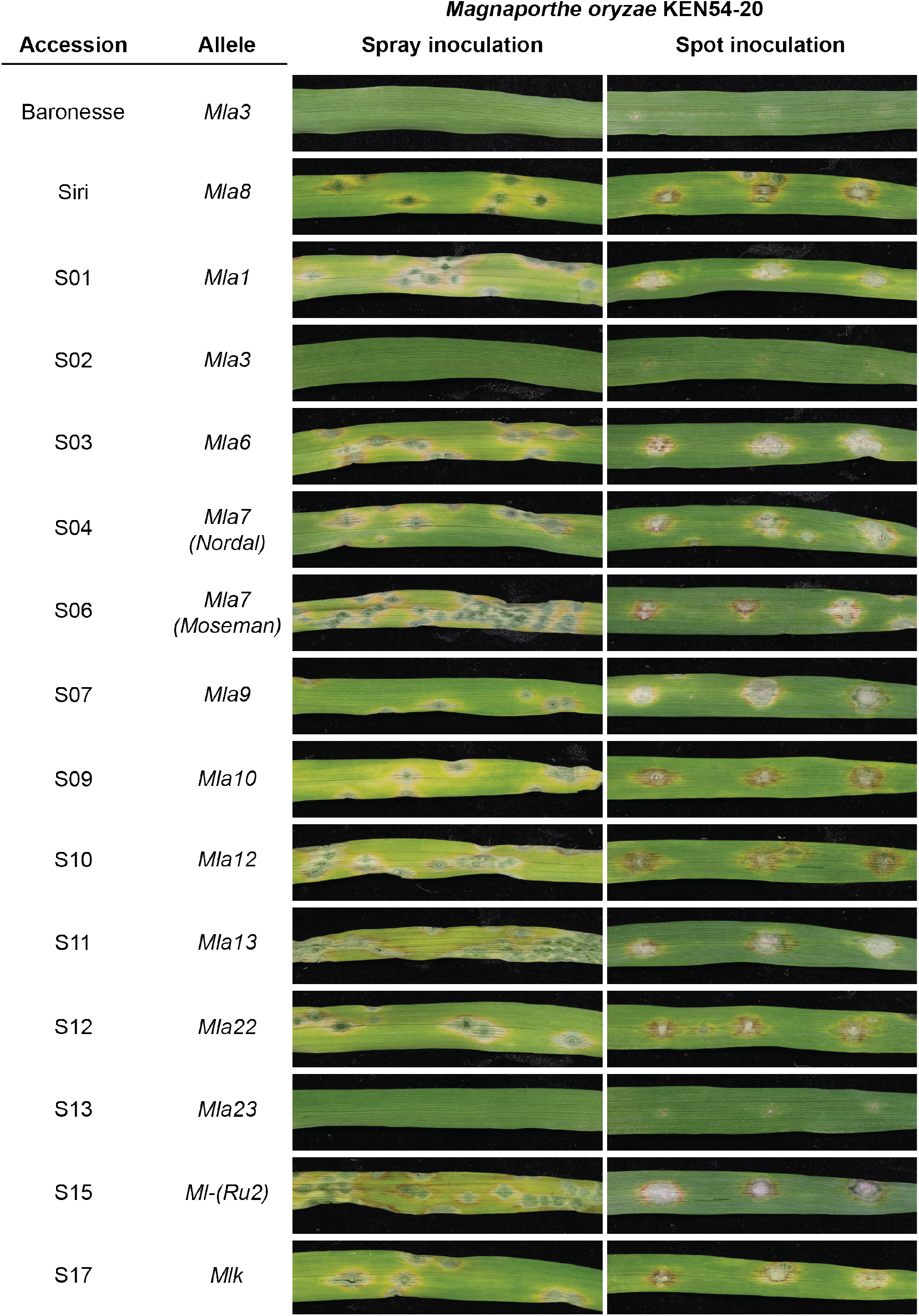
Barley accessions carrying *Mla3* and *Mla23* are resistant to *M. oryzae* KEN54-20. The barley powdery mildew Siri near-isogenic introgression line (NIL) panel were inoculated with *M. oryzae* KEN54-20 using spot and spray-based inoculation protocol. Controls include Baronesse (resistant) and Siri (susceptible). Siri NIL identifier and corresponding introgressed barley powdery mildew loci are indicated. Phenotypes are representative of three biological replicates with four technical replicates.

**Supplemental Figure 5.**
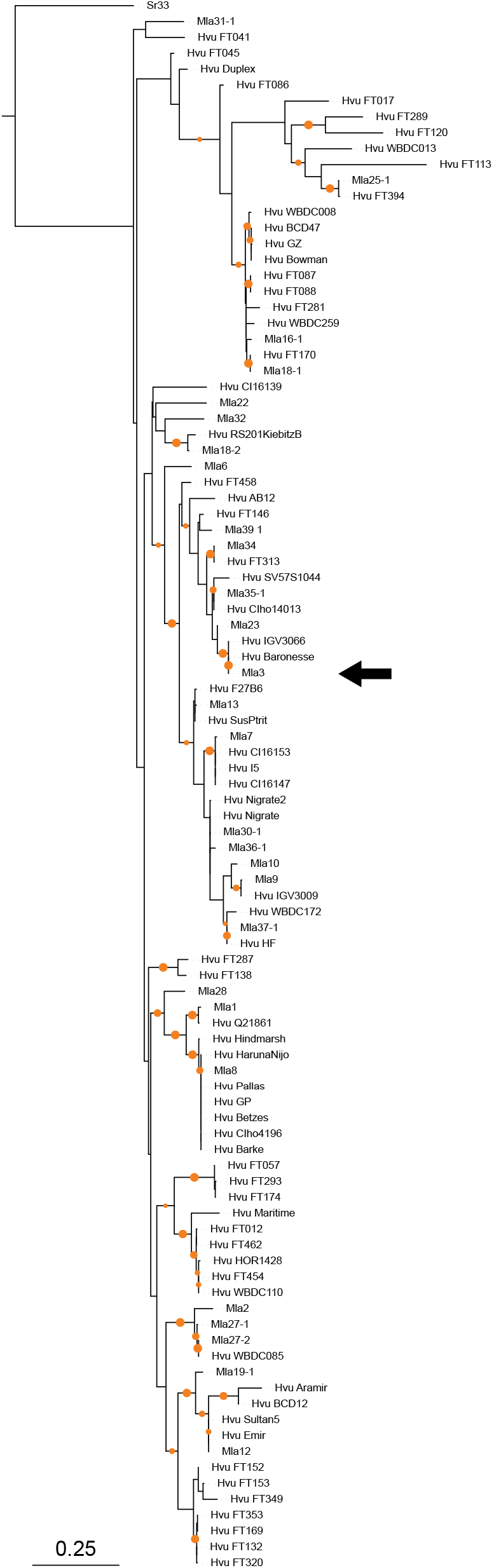
Mla3 and Mla23 are closely related RGH1 (Mla) alleles. Maximum likelihood phylogenetic tree of the RGH1 (Mla) family using protein sequence. Bootstrap support indicated by small and large orange dots indicate support from 80% to 90% and 90% to 100% based on 1,000 bootstraps, respectively. The position of Mla3 is indicated with an arrow.

**Supplemental Figure 6.**
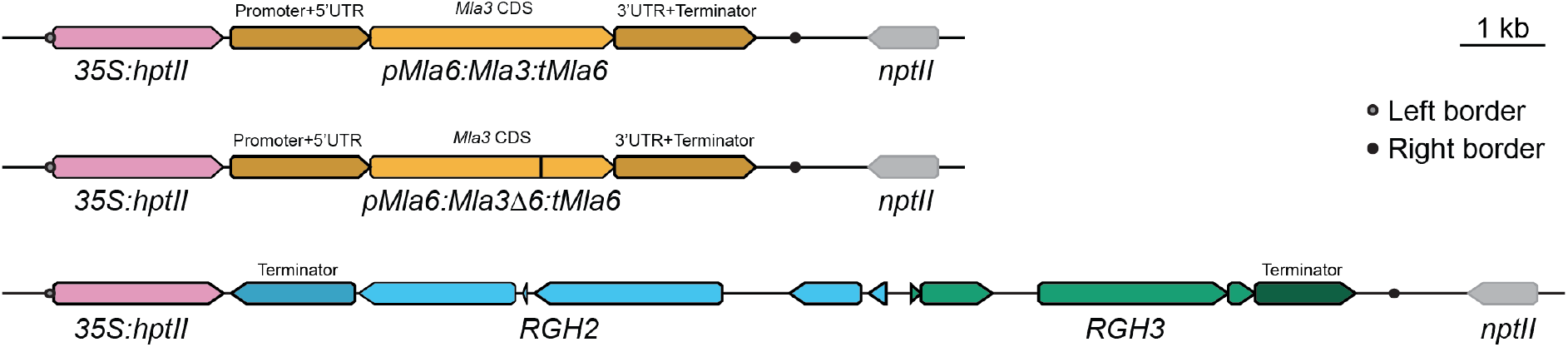
Transformation constructs for *Rmo1* candidate genes. The coding sequence of *Mla3* and *Mla3Δ6* were expressed under the promoter/5’UTR and 3’UTR/terminator of *Mla6*. *RGH2* and *RGH3* were maintained in their native genomic context in head-to-head orientation. All constructs were transformed into the pBract202 backbone that uses *nptII* as a bacterial selectable marker (shown in grey) and *hptII* driven by the 35S Cauliflower Mosaic Virus (35S) promoter for plant selection during transformation (shown in pink). Left and right T-DNA borders are shown with filled grey and black circles, respectively. Sequences of plasmids have been deposited in the NCBI database with accession codes OP561810 (*Mla3*), OP561809 (*Mla3△6*), and OP561811 (*RGH2*/*RGH3*).

**Supplemental Figure 7.**
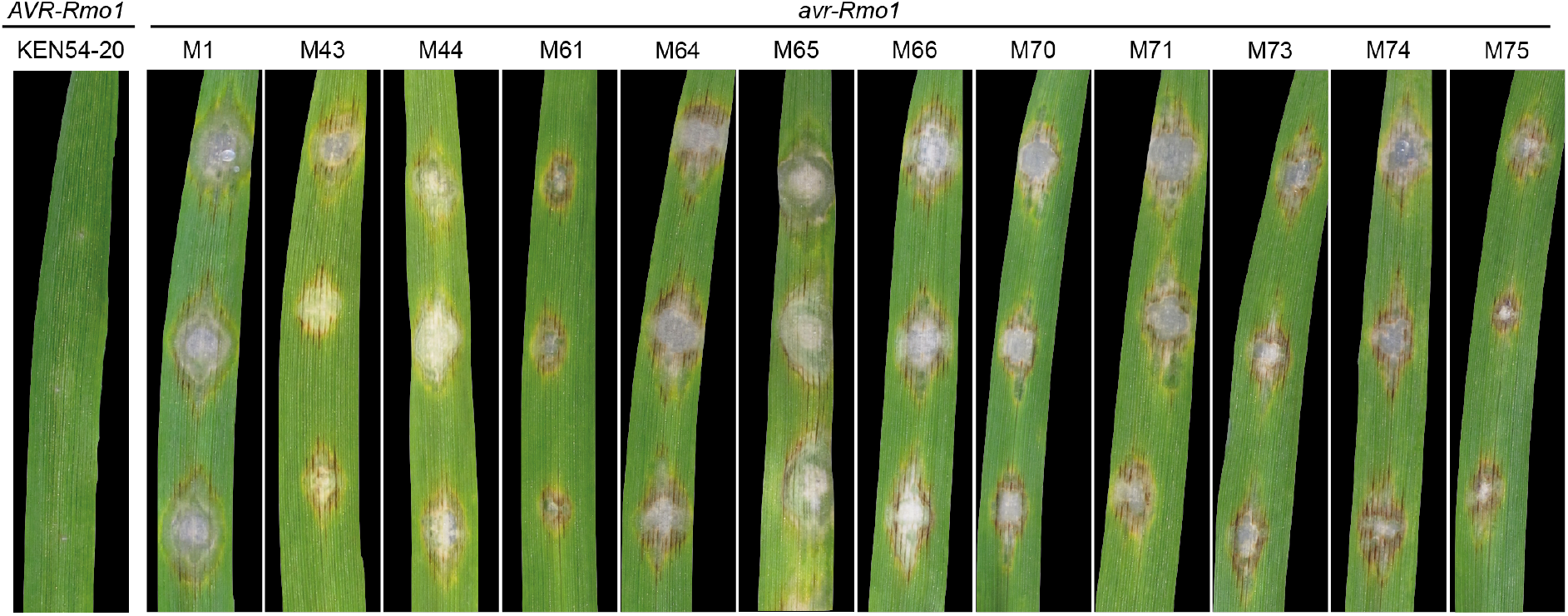
*avr-Rmo1* mutants are virulent on Baronesse. Baronesse (*Mla3*) leaves spot inoculated with *M. oryzae* KEN54-20 (*AVR-Rmo1*) and twelve independent *avr-Rmo1* mutants derived from UV mutagenesis. Phenotypes are representative of three biological replicates with three technical replicates.

**Supplemental Figure 8.**
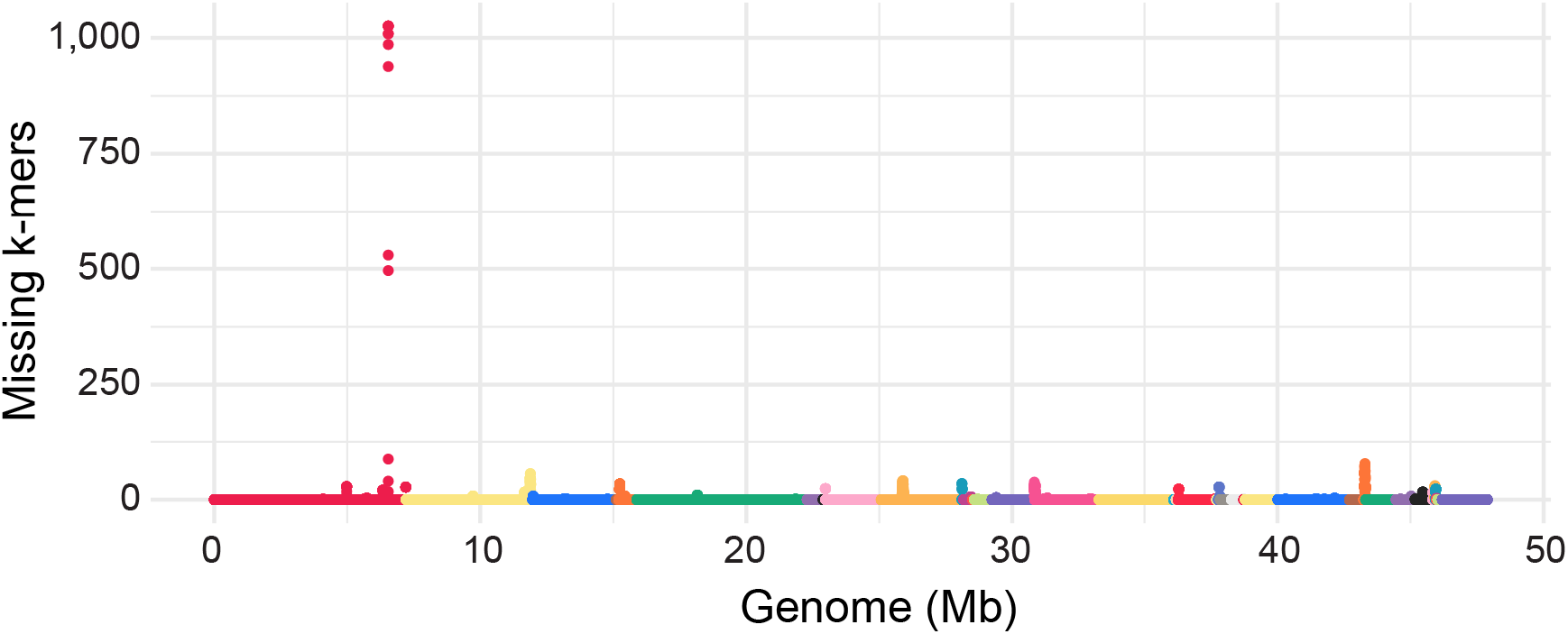
Genome-wide scan of regions with modified sequence relative to *M. oryzae* isolate Ken54-20. The number of shared missing *k*-mers among all mutants within a sliding window were determined using on window size of 10 kb and step size of 1 kb. Color coding corresponds to individual contigs of the Ken54-20 genome. The sharp peak corresponds to the *PWL2* locus on scaffold 183 (red).

**Supplemental Figure 9.**
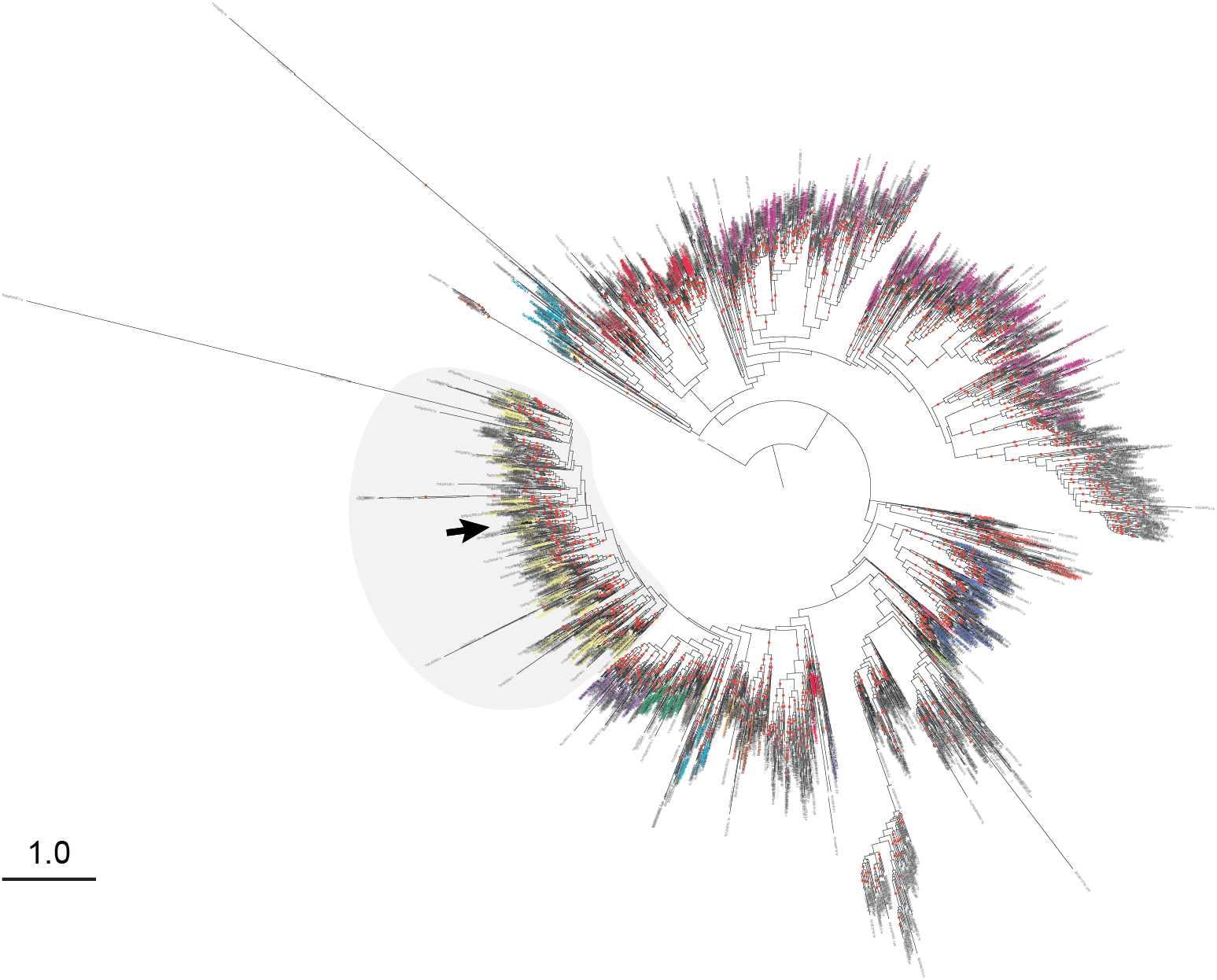
Structure-guided phylogenetic tree of NLR nucleotide binding domain from several PACMAD grasses. Maximum likelihood phylogenetic tree based on structure-guided multiple sequence alignment of the nucleotide binding domain (Pfam: PF00931) of NLRs from barley, rice, *B. distachyon*, *S. bicolor*, *S. italica*, maize, and weeping lovegrass. Color coding of clades based on original classification by Bailey *et al*. (2018) with the C17 clade in yellow. The position of *Mla* is indicated by an arrow. The NB domain from *Arabidopsis thaliana* ZAR1 (PDB: 6J5T) was used as outgroup for both phylogenetic trees. Dots on branches indicate bootstrap support exceeding 80%.

**Supplemental Figure 10.**
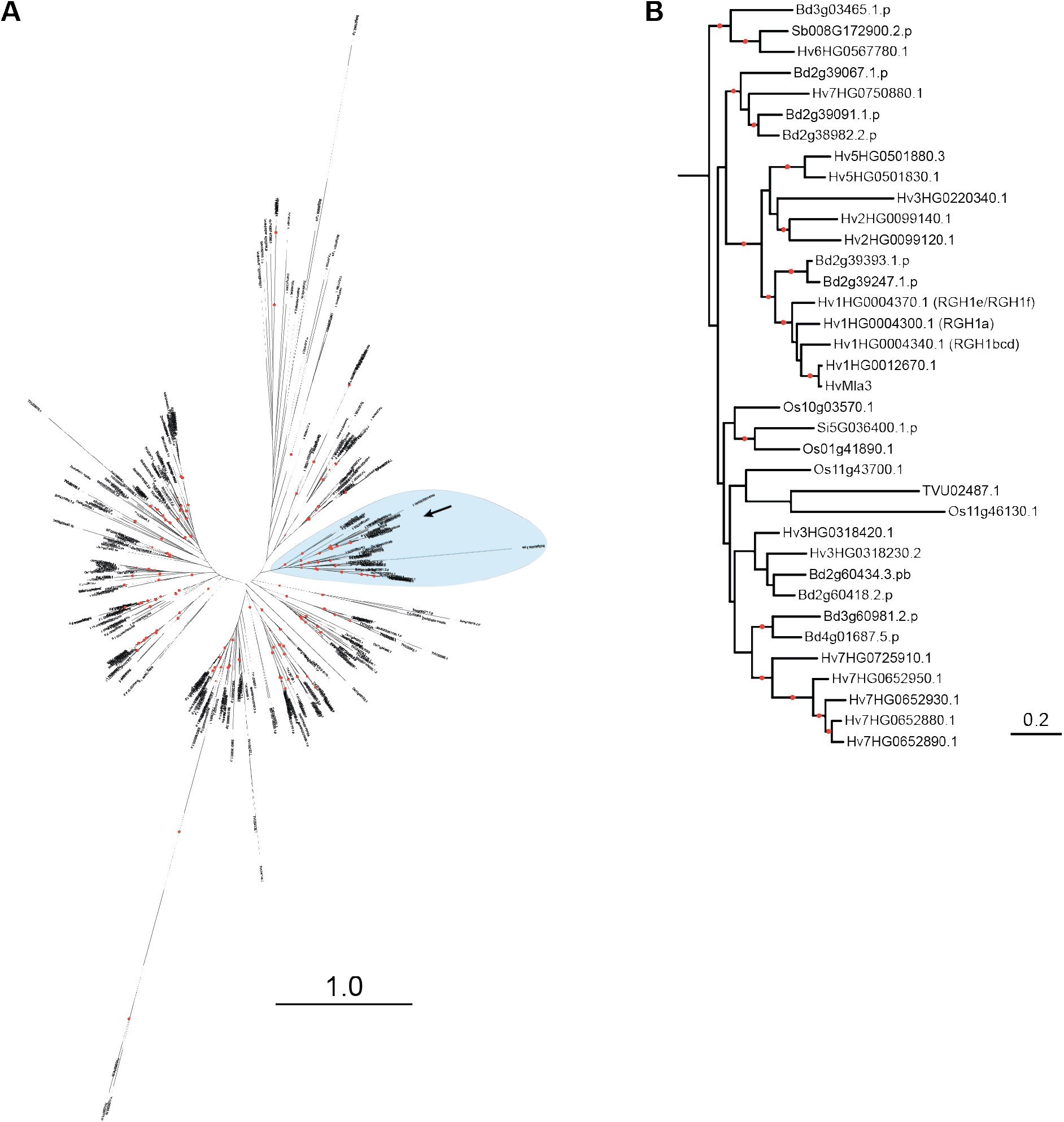
PACMAD grasses lack an ortholog of the RGH1 (*Mla*) gene family. **(A)** Maximum likelihood phylogenetic tree based on structure-guided multiple sequence alignment of the nucleotide binding domain of NLRs in the C17 clade from barley, rice, *B. distachyon*, *S. bicolor*, *S. italica*, maize, and weeping lovegrass. Light blue indicates the subclade containing the RGH1 (Mla) protein family. The position of *Mla* is indicated by an arrow. The NB domain from *Arabidopsis thaliana* ZAR1 (PDB: 6J5T) was used as outgroup. (B) The RGH1 gene family and related NLR sub-tree (in light blue) from **(A)**. The sub-tree contains NLRs from all grass species except maize. A single NLR from weeping lovegrass was found in the clade, TVU02487, which groups distinct from the RGH1 family. Bootstrap support with NLRs from both PACMAD and BOP grasses is observed within this sub-tree. For both trees, dots on branches indicate bootstrap support exceeding 80%.

**Supplementary Figure 11.**
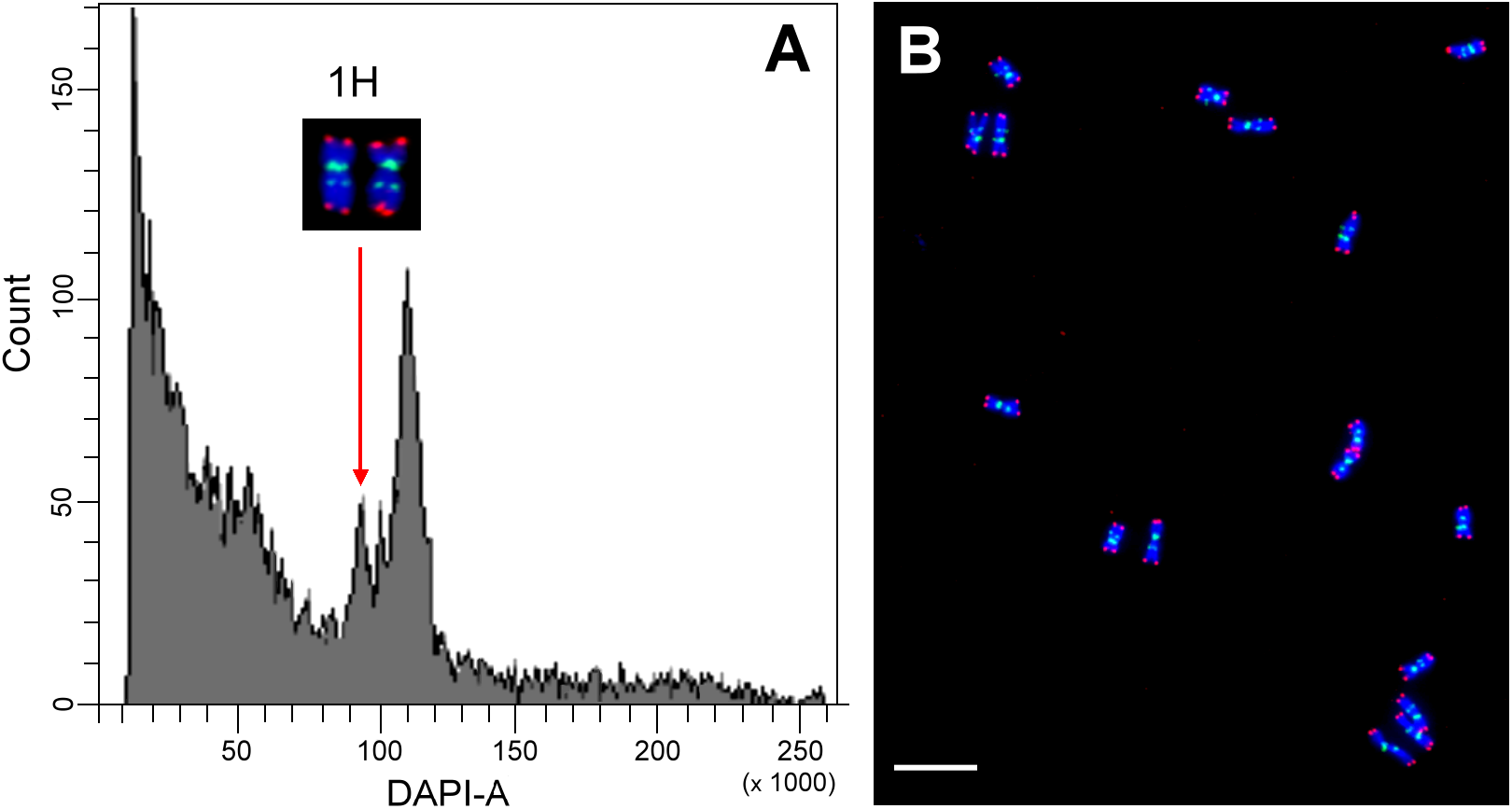
Flow cytometric analysis and sorting of barley cv. Baronesse chromosome 1H for long-range sequencing. **(A)** Flow cytometric analysis of DAPI-stained chromosomes resulted in histograms of relative fluorescence intensity (flow karyotypes) on which the peak representing chromosome 1H was clearly discriminated. This permitted sorting of chromosome 1H at purities ranging from 82.8 to 92.3%. **(B)** The purities in the sorted 1H fractions were determined by fluorescence *in situ* hybridization (FISH) using probes for GAA microsatellite (green) and HvT01 repeat (red) on chromosomes sorted onto microscope slides. The chromosomes were counterstained by DAPI (blue). Bar = 10 μm.

## Supplemental Data

**Supplemental Data 1. *De novo* assembled barley leaf transcriptomes.**

**Supplemental Data 2. Inventory of transgenic barley lines with *Mla3*, *Mla3△6*, *RGH2*/*RGH3*.**

**Supplemental Data 3. KASP markers used for fine-mapping of the *Mla3*/*Rmo1* locus.**

**Supplemental Data 4. Genomes used for NLR phylogenetic tree construction.**

## References

Ade, J., DeYoung, B. J., Golstein, C., & Innes, R. W. (2007). Indirect activation of a plant nucleotide binding site–leucine-rich repeat protein by a bacterial protease. Proceedings of the National Academy of Sciences, 104(7), 2531–2536.

Andrews, S. (2010). FastQC: a quality control tool for high throughput sequence data. In: Babraham Bioinformatics, Babraham Institute, Cambridge, United Kingdom.

Ashfield, T., Ong, L. E., Nobuta, K., Schneider, C. M., & Innes, R. W. (2004). Convergent evolution of disease resistance gene specificity in two flowering plant families. The Plant Cell, 16(2), 309–318.

Ashfield, T., Redditt, T., Russell, A., Kessens, R., Rodibaugh, N., Galloway, L., Kang, Q., Podicheti, R., & Innes, R. W. (2014). Evolutionary relationship of disease resistance genes in soybean and Arabidopsis specific for the Pseudomonas syringae effectors AvrB and AvrRpm1. Plant physiology, 166(1), 235–251.

Axtell, M. J., & Staskawicz, B. J. (2003). Initiation of RPS2-specified disease resistance in Arabidopsis is coupled to the AvrRpt2-directed elimination of RIN4. Cell, 112(3), 369–377.

Bailey, P. C., Schudoma, C., Jackson, W., Baggs, E., Dagdas, G., Haerty, W., Moscou, M., & Krasileva, K. V. (2018). Dominant integration locus drives continuous diversification of plant immune receptors with exogenous domain fusions. Genome biology, 19(1), 1–18.

Baudin, M., Hassan, J. A., Schreiber, K. J., & Lewis, J. D. (2017). Analysis of the ZAR1 immune complex reveals determinants for immunity and molecular interactions. Plant physiology, 174(4), 2038–2053.

Bauer, S., Yu, D., Lawson, A. W., Saur, I. M., Frantzeskakis, L., Kracher, B., Logemann, E., Chai, J., Maekawa, T., & Schulze-Lefert, P. (2021). The leucine-rich repeats in allelic barley MLA immune receptors define specificity towards sequence-unrelated powdery mildew avirulence effectors with a predicted common RNase-like fold. PLoS pathogens, 17(2), e1009223.

Bennetzen, J. L., Schmutz, J., Wang, H., Percifield, R., Hawkins, J., Pontaroli, A. C., Estep, M., Feng, L., Vaughn, J. N., Grimwood, J., Jenkins, J., Barry, K., Lindquist, E., Hellsten, U., Deshpande, S., Wang, X., Wu, X., Mitros, T., Triplett, J.,. .. Devos, K. M. (2012). Reference genome sequence of the model plant Setaria. Nature Biotechnology, 30(6), 555–561. https://doi.org/10.1038/nbt.2196

Bettgenhaeuser, J., Hernández-Pinzón, I., Dawson, A. M., Gardiner, M., Green, P., Taylor, J., Smoker, M., Ferguson, J. N., Emmrich, P., & Hubbard, A. (2021). The barley immune receptor Mla recognizes multiple pathogens and contributes to host range dynamics. Nature communications, 12(1), 1–14.

Bilgic, H., Steffenson, B., & Hayes, P. (2006). Molecular mapping of loci conferring resistance to different pathotypes of the spot blotch pathogen in barley. Phytopathology, 96(7), 699–708.

Bolger, A. M., Lohse, M., & Usadel, B. (2014). Trimmomatic: a flexible trimmer for Illumina sequence data. Bioinformatics, 30(15), 2114–2120.

Bourras, S., Kunz, L., Xue, M., Praz, C. R., Müller, M. C., Kälin, C., Schläfli, M., Ackermann, P., Flückiger, S., & Parlange, F. (2019). The AvrPm3-Pm3 effector-NLR interactions control both race-specific resistance and host-specificity of cereal mildews on wheat. Nature communications, 10(1), 1–16.

Bourras, S., Praz, C. R., Spanu, P. D., & Keller, B. (2018). Cereal powdery mildew effectors: a complex toolbox for an obligate pathogen. Current opinion in microbiology, 46, 26–33.

Brabham, H. J., Hernández-Pinzón, I., Holden, S., Lorang, J., & Moscou, M. J. (2018). An ancient integration in a plant NLR is maintained as a trans-species polymorphism. BioRxiv, 239541.

Briggs, F. N., & Stanford, E. H. (1938). Linkage of factors for resistance to mildew in barley. Journal of Genetics, 37(1), 107–117.

Broman, K. W., Wu, H., Sen, Ś., & Churchill, G. A. (2003). R/qtl: QTL mapping in experimental crosses. bioinformatics, 19(7), 889–890.

Brown, J. K., & Tellier, A. (2011). Plant-parasite coevolution: bridging the gap between genetics and ecology. Annual review of phytopathology, 49(1), 345–367.

Caldwell, K. S., & Michelmore, R. W. (2009). Arabidopsis thaliana genes encoding defense signaling and recognition proteins exhibit contrasting evolutionary dynamics. Genetics, 181(2), 671–684.

Camacho, C., Coulouris, G., Avagyan, V., Ma, N., Papadopoulos, J., Bealer, K., & Madden, T. L. (2009). BLAST+: architecture and applications. BMC bioinformatics, 10(1), 1–9.

Carballo, J., Santos, B. A. C. M., Zappacosta, D., Garbus, I., Selva, J. P., Gallo, C. A., Díaz, A., Albertini, E., Caccamo, M., & Echenique, V. (2019). A high-quality genome of Eragrostis curvula grass provides insights into Poaceae evolution and supports new strategies to enhance forage quality. Scientific Reports, 9(1), 10250. https://doi.org/10.1038/s41598-019-46610-0

Carter, M. E., Helm, M., Chapman, A. V., Wan, E., Restrepo Sierra, A. M., Innes, R. W., Bogdanove, A. J., & Wise, R. P. (2019). Convergent evolution of effector protease recognition by Arabidopsis and barley. Molecular Plant-Microbe Interactions, 32(5), 550–565.

Cesari, S. (2018). Multiple strategies for pathogen perception by plant immune receptors. New Phytol, 219(1), 17–24. https://doi.org/10.1111/nph.14877

Cesari, S., Bernoux, M., Moncuquet, P., Kroj, T., & Dodds, P. N. (2014). A novel conserved mechanism for plant NLR protein pairs: the “integrated decoy” hypothesis. Front Plant Sci, 5, 606. https://doi.org/10.3389/fpls.2014.00606

Chen, J., Upadhyaya, N. M., Ortiz, D., Sperschneider, J., Li, F., Bouton, C., Breen, S., Dong, C., Xu, B., & Zhang, X. (2017). Loss of AvrSr50 by somatic exchange in stem rust leads to virulence for Sr50 resistance in wheat. Science, 358(6370), 1607–1610.

Close, T. J., Bhat, P. R., Lonardi, S., Wu, Y., Rostoks, N., Ramsay, L., Druka, A., Stein, N., Svensson, J. T., & Wanamaker, S. (2009). Development and implementation of high-throughput SNP genotyping in barley. BMC genomics, 10(1), 1–13.

Cooley, M. B., Pathirana, S., Wu, H.-J., Kachroo, P., & Klessig, D. F. (2000). Members of the Arabidopsis HRT/RPP8 family of resistance genes confer resistance to both viral and oomycete pathogens. The Plant Cell, 12(5), 663–676.

Couch, B. C., & Kohn, L. M. (2002). A multilocus gene genealogy concordant with host preference indicates segregation of a new species, Magnaporthe oryzae, from M. grisea. Mycologia, 94(4), 683–693.

Dangl, J. L., & Jones, J. D. (2001). Plant pathogens and integrated defence responses to infection. Nature, 411(6839), 826–833.

Dawson, A. M., Ferguson, J. N., Gardiner, M., Green, P., Hubbard, A., & Moscou, M. J. (2016). Isolation and fine mapping of Rps6: an intermediate host resistance gene in barley to wheat stripe rust. Theoretical and applied genetics, 129(4), 831–843.

de Guillen, K., Ortiz-Vallejo, D., Gracy, J., Fournier, E., Kroj, T., & Padilla, A. (2015). Structure analysis uncovers a highly diverse but structurally conserved effector family in phytopathogenic fungi. PLoS pathogens, 11(10), e1005228.

Deslandes, L., Olivier, J., Peeters, N., Feng, D. X., Khounlotham, M., Boucher, C., Somssich, I., Genin, S., & Marco, Y. (2003). Physical interaction between RRS1-R, a protein conferring resistance to bacterial wilt, and PopP2, a type III effector targeted to the plant nucleus. Proceedings of the National Academy of Sciences, 100(13), 8024–8029.

DeYoung, B. J., Qi, D., Kim, S. H., Burke, T. P., & Innes, R. W. (2012). Activation of a plant nucleotide binding-leucine rich repeat disease resistance protein by a modified self protein. Cellular microbiology, 14(7), 1071–1084.

Ding, J., Zhang, W., Jing, Z., Chen, J.-Q., & Tian, D. (2007). Unique pattern of R-gene variation within populations in Arabidopsis. Molecular Genetics and Genomics, 277(6), 619–629.

Doležel, J., Lucretti, S., Molnár, I., Cápal, P., Giorgi, D. (2021). Chromosome analysis and sorting. Cytometry 99(4), 328–342.

Flor, H. (1956). The complementary genic systems in flax and flax rust. Advances in genetics, 8, 29–54.

Franceschetti, M., Maqbool, A., Jimenez-Dalmaroni, M. J., Pennington, H. G., Kamoun, S., & Banfield, M. J. (2017). Effectors of Filamentous Plant Pathogens: Commonalities amid Diversity. Microbiol Mol Biol Rev, 81(2). https://doi.org/10.1128/MMBR.00066-16

Fujisaki, K., Abe, Y., Ito, A., Saitoh, H., Yoshida, K., Kanzaki, H., Kanzaki, E., Utsushi, H., Yamashita, T., & Kamoun, S. (2015). Rice Exo70 interacts with a fungal effector, AVR-Pii, and is required for AVR-Pii-triggered immunity. The Plant Journal, 83(5), 875–887.

Gassmann, W., Hinsch, M. E., & Staskawicz, B. J. (1999). The Arabidopsis RPS4 bacterial-resistance gene is a member of the TIR-NBS-LRR family of disease-resistance genes. The Plant Journal, 20(3), 265–277.

Gibson, D. G., Young, L., Chuang, R.-Y., Venter, J. C., Hutchison, C. A., & Smith, H. O. (2009). Enzymatic assembly of DNA molecules up to several hundred kilobases. Nature methods, 6(5), 343–345.

Gladieux, P., Condon, B., Ravel, S., Soanes, D., Maciel, J. L. N., Nhani Jr, A., Chen, L., Terauchi, R., Lebrun, M.-H., & Tharreau, D. (2018). Gene flow between divergent cereal-and grass-specific lineages of the rice blast fungus Magnaporthe oryzae. MBio, 9(1), e01219–01217.

Goff, S. A., Ricke, D., Lan, T.-H., Presting, G., Wang, R., Dunn, M., Glazebrook, J., Sessions, A., Oeller, P., Varma, H., Hadley, D., Hutchison, D., Martin, C., Katagiri, F., Lange, B. M., Moughamer, T., Xia, Y., Budworth, P., Zhong, J.,. .. Briggs, S. (2002). A Draft Sequence of the Rice Genome (*Oryza sativa* L. ssp. *japonica*). Science, 296(5565), 92–100. https://doi.org/doi:10.1126/science.1068275

Goggin, F. L., Jia, L., Shah, G., Hebert, S., Williamson, V. M., & Ullman, D. E. (2006). Heterologous expression of the Mi-1.2 gene from tomato confers resistance against nematodes but not aphids in eggplant. Molecular Plant-Microbe Interactions, 19(4), 383–388.

Grabherr, M. G., Haas, B. J., Yassour, M., Levin, J. Z., Thompson, D. A., Amit, I., Adiconis, X., Fan, L., Raychowdhury, R., & Zeng, Q. (2011). Trinity: reconstructing a full-length transcriptome without a genome from RNA-Seq data. Nature biotechnology, 29(7), 644.

Halterman, D., Zhou, F., Wei, F., Wise, R. P., & Schulze-Lefert, P. (2001). The MLA6 coiled-coil, NBS-LRR protein confers AvrMla6-dependent resistance specificity to Blumeria graminis f. sp. hordei in barley and wheat. The Plant Journal, 25(3), 335–348.

Hensel, G., Kastner, C., Oleszczuk, S., Riechen, J., & Kumlehn, J. (2009). Agrobacterium-mediated gene transfer to cereal crop plants: current protocols for barley, wheat, triticale, and maize. International Journal of Plant Genomics, 2009.

Hensel, G., & Kumlehn, J. (2004). Genetic transformation of barley (Hordeum vulgare L.) by co-culture of immature embryos with Agrobacterium. In Transgenic Crops of the World (pp. 35–44). Springer.

Holden, S., Bergum, M., Green, P., Bettgenhaeuser, J., Hernández-Pinzón, I., Thind, A., Clare, S., Russell, J. M., Hubbard, A., & Taylor, J. (2022). A lineage-specific Exo70 is required for receptor kinase–mediated immunity in barley. Science Advances, 8(27), eabn7258.

Holm, L. (2022). Dali server: structural unification of protein families. Nucleic Acids Research.

Huang, J., Si, W., Deng, Q., Li, P., & Yang, S. (2014). Rapid evolution of avirulence genes in rice blast fungus Magnaporthe oryzae. BMC genetics, 15(1), 1–10.

Inoue, Y., Vy, T. T., Yoshida, K., Asano, H., Mitsuoka, C., Asuke, S., Anh, V. L., Cumagun, C. J., Chuma, I., & Terauchi, R. (2017). Evolution of the wheat blast fungus through functional losses in a host specificity determinant. Science, 357(6346), 80–83.

Inukai, T., Vales, M. I., Hori, K., Sato, K., & Hayes, P. M. (2006). RMo 1 confers blast resistance in barley and is located within the complex of resistance genes containing Mla, a powdery mildew resistance gene. Molecular plant-microbe interactions, 19(9), 1034–1041.

Jacob, S. (2021). Magnaporthe Oryzae: Methods and Protocols. Springer.

Jia, Y., McAdams, S. A., Bryan, G. T., Hershey, H. P., & Valent, B. (2000). Direct interaction of resistance gene and avirulence gene products confers rice blast resistance. The EMBO journal, 19(15), 4004–4014.

Jia, Y., Valent, B., & Lee, F. (2003). Determination of host responses to Magnaporthe grisea on detached rice leaves using a spot inoculation method. Plant Disease, 87(2), 129–133.

Jones, J. D., & Dangl, J. L. (2006). The plant immune system. Nature, 444(7117), 323–329. https://doi.org/10.1038/nature05286

Jørgensen, J. H., & Wolfe, M. (1994). Genetics of powdery mildew resistance in barley. Critical Reviews in Plant Sciences, 13(1), 97–119.

Jumper, J., Evans, R., Pritzel, A., Green, T., Figurnov, M., Ronneberger, O., Tunyasuvunakool, K., Bates, R., Žídek, A., & Potapenko, A. (2021). Highly accurate protein structure prediction with AlphaFold. Nature, 596(7873), 583–589.

Kang, S., Sweigard, J. A., & Valent, B. (1995). The PWL host specificity gene family in the blast fungus Magnaporthe grisea. MPMI-Molecular Plant Microbe Interactions, 8(6), 939–948.

Khang, C. H., Berruyer, R., Giraldo, M. C., Kankanala, P., Park, S.-Y., Czymmek, K., Kang, S., & Valent, B. (2010). Translocation of Magnaporthe oryzae effectors into rice cells and their subsequent cell-to-cell movement. The Plant Cell, 22(4), 1388–1403.

Kim, H., Prokchorchik, M., & Sohn, K. H. (2022). Investigation of natural <scp>RIN4</scp> variants reveals a motif crucial for function and provides an opportunity to broaden <scp>NLR</scp> regulation specificity. The Plant Journal, 110(1), 58–70. https://doi.org/10.1111/tpj.15653

Kim, H.-S., Desveaux, D., Singer, A. U., Patel, P., Sondek, J., & Dangl, J. L. (2005). The Pseudomonas syringae effector AvrRpt2 cleaves its C-terminally acylated target, RIN4, from Arabidopsis membranes to block RPM1 activation. Proceedings of the National Academy of Sciences, 102(18), 6496–6501.

Kim, M. G., Geng, X., Lee, S. Y., & Mackey, D. (2009). The Pseudomonas syringae type III effector AvrRpm1 induces significant defenses by activating the Arabidopsis nucleotide-binding leucine-rich repeat protein RPS2. The Plant Journal, 57(4), 645–653.

Kim, S. H., Qi, D., Ashfield, T., Helm, M., & Innes, R. W. (2016). Using decoys to expand the recognition specificity of a plant disease resistance protein. Science, 351(6274), 684–687.

Kinizios, S., Jahoor, A., & Fischbeck, G. (1995). Powdery-mildew-resistance genes Mla29 and Mla32 in H. spontaneum derived winter-barley lines. Plant Breeding, 114(3), 265–266.

Kølster, P., & Stølen, O. (1987). Barley isolines with genes for resistance to Erysiphe graminis f. sp. hordei in the recurrent parent ‘Siri’. Plant breeding, 98(1), 79–82.

Kourelis, J., & van der Hoorn, R. A. L. (2018). Defended to the Nines: 25 Years of Resistance Gene Cloning Identifies Nine Mechanisms for R Protein Function. Plant Cell, 30(2), 285–299. https://doi.org/10.1105/tpc.17.00579

Kuang, H., Caldwell, K. S., Meyers, B. C., & Michelmore, R. W. (2008). Frequent sequence exchanges between homologs of RPP8 in Arabidopsis are not necessarily associated with genomic proximity. The Plant Journal, 54(1), 69–80.

Lai, Y., & Eulgem, T. (2018). Transcript-level expression control of plant NLR genes. Molecular plant pathology, 19(5), 1267–1281.

Le Roux, C., Huet, G., Jauneau, A., Camborde, L., Trémousaygue, D., Kraut, A., Zhou, B., Levaillant, M., Adachi, H., & Yoshioka, H. (2015). A receptor pair with an integrated decoy converts pathogen disabling of transcription factors to immunity. Cell, 161(5), 1074–1088.

Leng, Y., Zhao, M., Fiedler, J., Dreiseitl, A., Chao, S., Li, X., & Zhong, S. (2020). Molecular mapping of loci conferring susceptibility to spot blotch and resistance to powdery mildew in barley using the sequencing-based genotyping approach. Phytopathology, 110(2), 440–446.

Leng, Y., Zhao, M., Wang, R., Steffenson, B. J., Brueggeman, R. S., & Zhong, S. (2018). The gene conferring susceptibility to spot blotch caused by Cochliobolus sativus is located at the Mla locus in barley cultivar Bowman. Theoretical and Applied Genetics, 131(7), 1531–1539.

Lewis, J. D., Lee, A. H.-Y., Hassan, J. A., Wan, J., Hurley, B., Jhingree, J. R., Wang, P. W., Lo, T., Youn, J.-Y., & Guttman, D. S. (2013). The Arabidopsis ZED1 pseudokinase is required for ZAR1-mediated immunity induced by the Pseudomonas syringae type III effector HopZ1a. Proceedings of the National Academy of Sciences, 110(46), 18722–18727.

Li, H., Handsaker, B., Wysoker, A., Fennell, T., Ruan, J., Homer, N., Marth, G., Abecasis, G., & Durbin, R. (2009). The sequence alignment/map format and SAMtools. Bioinformatics, 25(16), 2078–2079.

Li, Z., Huang, J., Wang, Z., Meng, F., Zhang, S., Wu, X., Zhang, Z., & Gao, Z. (2019). Overexpression of Arabidopsis Nucleotide-Binding and Leucine-Rich Repeat Genes RPS2 and RPM1(D505V) Confers Broad-Spectrum Disease Resistance in Rice. Front Plant Sci, 10, 417. https://doi.org/10.3389/fpls.2019.00417

Lorang, J., Cuesta-Marcos, A., Hayes, P., & Wolpert, T. (2010). Identification and mapping of adult-onset sensitivity to victorin in barley. Molecular breeding, 26(3), 545–550.

Lorang, J., Hagerty, C., Lee, R., McClean, P., & Wolpert, T. (2018). Genetic analysis of victorin sensitivity and identification of a causal nucleotide-binding site leucine-rich repeat gene in Phaseolus vulgaris. Molecular Plant-Microbe Interactions, 31(10), 1069–1074.

Lorang, J. M., Carkaci-Salli, N., & Wolpert, T. J. (2004). Identification and characterization of victorin sensitivity in Arabidopsis thaliana. Molecular plant-microbe interactions, 17(6), 577–582.

Lorang, J. M., Sweat, T. A., & Wolpert, T. J. (2007). Plant disease susceptibility conferred by a “resistance” gene. Proceedings of the National Academy of Sciences, 104(37), 14861–14866.

Lu, X., Kracher, B., Saur, I. M., Bauer, S., Ellwood, S. R., Wise, R., Yaeno, T., Maekawa, T., & Schulze-Lefert, P. (2016). Allelic barley MLA immune receptors recognize sequence-unrelated avirulence effectors of the powdery mildew pathogen. Proceedings of the National Academy of Sciences, 113(42), E6486–E6495.

Ma, Y., Guo, H., Hu, L., Martinez, P. P., Moschou, P. N., Cevik, V., Ding, P., Duxbury, Z., Sarris, P. F., & Jones, J. D. (2018). Distinct modes of derepression of an Arabidopsis immune receptor complex by two different bacterial effectors. Proceedings of the National Academy of Sciences, 115(41), 10218–10227.

Mackey, D., Belkhadir, Y., Alonso, J. M., Ecker, J. R., & Dangl, J. L. (2003). Arabidopsis RIN4 is a target of the type III virulence effector AvrRpt2 and modulates RPS2-mediated resistance. Cell, 112(3), 379–389.

Mackey, D., Holt III, B. F., Wiig, A., & Dangl, J. L. (2002). RIN4 interacts with Pseudomonas syringae type III effector molecules and is required for RPM1-mediated resistance in Arabidopsis. Cell, 108(6), 743–754.

Maekawa, T., Kracher, B., Saur, I. M., Yoshikawa-Maekawa, M., Kellner, R., Pankin, A., von Korff, M., & Schulze-Lefert, P. (2019). Subfamily-specific specialization of RGH1/MLA immune receptors in wild barley. Molecular Plant-Microbe Interactions, 32(1), 107–119.

Mapleson, D., Garcia Accinelli, G., Kettleborough, G., Wright, J., & Clavijo, B. J. (2017). KAT: a K-mer analysis toolkit to quality control NGS datasets and genome assemblies. Bioinformatics, 33(4), 574–576.

Märkle, H., Saur, Isabel M. L., & Stam, R. (2022). Evolution of resistance (R) gene specificity. Essays in Biochemistry. https://doi.org/10.1042/ebc20210077

Mascher, M., Wicker, T., Jenkins, J., Plott, C., Lux, T., Koh, C. S., Ens, J., Gundlach, H., Boston, L. B., & Tulpová, Z. (2021). Long-read sequence assembly: a technical evaluation in barley. The Plant Cell, 33(6), 1888–1906.

Mazo-Molina, C., Mainiero, S., Haefner, B. J., Bednarek, R., Zhang, J., Feder, A., Shi, K., Strickler, S. R., & Martin, G. B. (2020). Ptr1 evolved convergently with RPS2 and Mr5 to mediate recognition of AvrRpt2 in diverse solanaceous species. The Plant Journal, 103(4), 1433–1445. https://doi.org/https://doi.org/10.1111/tpj.14810

Mazo-Molina, C., Mainiero, S., Hind, S. R., Kraus, C. M., Vachev, M., Maviane-Macia, F., Lindeberg, M., Saha, S., Strickler, S. R., & Feder, A. (2019). The Ptr1 locus of Solanum lycopersicoides confers resistance to race 1 strains of Pseudomonas syringae pv. tomato and to Ralstonia pseudosolanacearum by recognizing the type III effectors AvrRpt2 and RipBN. Molecular Plant-Microbe Interactions, 32(8), 949–960.

McDowell, J. M., Dhandaydham, M., Long, T. A., Aarts, M. G., Goff, S., Holub, E. B., & Dangl, J. L. (1998). Intragenic recombination and diversifying selection contribute to the evolution of downy mildew resistance at the RPP8 locus of Arabidopsis. The Plant Cell, 10(11), 1861–1874.

Mukhi, N., Brown, H., Gorenkin, D., Ding, P., Bentham, A. R., Stevenson, C. E., Jones, J. D., & Banfield, M. J. (2021). Perception of structurally distinct effectors by the integrated WRKY domain of a plant immune receptor. Proceedings of the National Academy of Sciences, 118(50), e2113996118.

Muñoz-Amatriaín, M., Moscou, M. J., Bhat, P. R., Svensson, J. T., Bartoš, J., Suchánková, P., Šimková, H., Endo, T. R., Fenton, R. D., & Lonardi, S. (2011). An improved consensus linkage map of barley based on flow-sorted chromosomes and single nucleotide polymorphism markers. The Plant Genome, 4(3).

Narusaka, M., Iuchi, S., & Narusaka, Y. (2017). Analyses of natural variation indicates that the absence of RPS4/RRS1 and amino acid change in RPS4 cause loss of their functions and resistance to pathogens. Plant signaling & behavior, 12(3), e1293218.

Narusaka, M., Kubo, Y., Hatakeyama, K., Imamura, J., Ezura, H., Nanasato, Y., Tabei, Y., Takano, Y., Shirasu, K., & Narusaka, Y. (2013). Interfamily transfer of dual NB-LRR genes confers resistance to multiple pathogens. PLoS One, 8(2), e55954.

Narusaka, M., Shirasu, K., Noutoshi, Y., Kubo, Y., Shiraishi, T., Iwabuchi, M., & Narusaka, Y. (2009). RRS1 and RPS4 provide a dual Resistance-gene system against fungal and bacterial pathogens. The Plant Journal, 60(2), 218–226.

Ngou, B. P. M., Ding, P., & Jones, J. D. G. (2022). Thirty years of resistance: Zig-zag through the plant immune system. Plant Cell, 34(5), 1447–1478. https://doi.org/10.1093/plcell/koac041

Nombela, G., Williamson, V. M., & Muñiz, M. (2003). The root-knot nematode resistance gene Mi-1.2 of tomato is responsible for resistance against the whitefly Bemisia tabaci. Molecular Plant-Microbe Interactions, 16(7), 645–649.

Ou, S. H. (1985). Rice diseases (2nd ed. ed.). Slough: CAB Internationl, 1985.

Parker, D., Beckmann, M., Enot, D. P., Overy, D. P., Rios, Z. C., Gilbert, M., Talbot, N., & Draper, J. (2008). Rice blast infection of Brachypodium distachyon as a model system to study dynamic host/pathogen interactions. Nature Protocols, 3(3), 435–445.

Paterson, A. H., Bowers, J. E., Bruggmann, R., Dubchak, I., Grimwood, J., Gundlach, H., Haberer, G., Hellsten, U., Mitros, T., Poliakov, A., Schmutz, J., Spannagl, M., Tang, H., Wang, X., Wicker, T., Bharti, A. K., Chapman, J., Feltus, F. A., Gowik, U.,. .. Rokhsar, D. S. (2009). The Sorghum bicolor genome and the diversification of grasses. Nature, 457(7229), 551–556. https://doi.org/10.1038/nature07723

Periyannan, S., Moore, J., Ayliffe, M., Bansal, U., Wang, X., Huang, L., Deal, K., Luo, M., Kong, X., & Bariana, H. (2013). The gene Sr33, an ortholog of barley Mla genes, encodes resistance to wheat stem rust race Ug99. Science, 341(6147), 786–788.

Prokchorchik, M., Choi, S., Chung, E.-H., Won, K., Dangl, J. L., & Sohn, K. H. (2020). A host target of a bacterial cysteine protease virulence effector plays a key role in convergent evolution of plant innate immune system receptors. New Phytologist, 225(3), 1327–1342. https://doi.org/https://doi.org/10.1111/nph.16218

Qi, D., Dubiella, U., Kim, S. H., Sloss, D. I., Dowen, R. H., Dixon, J. E., & Innes, R. W. (2014). Recognition of the protein kinase AVRPPHB SUSCEPTIBLE1 by the disease resistance protein RESISTANCE TO PSEUDOMONAS SYRINGAE5 is dependent on s-acylation and an exposed loop in AVRPPHB SUSCEPTIBLE1. Plant physiology, 164(1), 340–351.

Ravensdale, M., Nemri, A., Thrall, P. H., Ellis, J. G., & Dodds, P. N. (2011). Co-evolutionary interactions between host resistance and pathogen effector genes in flax rust disease. Molecular plant pathology, 12(1), 93–102.

Robinson, J. T., Thorvaldsdóttir, H., Winckler, W., Guttman, M., Lander, E. S., Getz, G., & Mesirov, J. P. (2011). Integrative genomics viewer. Nat Biotechnol, 29(1), 24–26. https://doi.org/10.1038/nbt.1754

Russell, A. R., Ashfield, T., & Innes, R. W. (2015). Pseudomonas syringae effector AvrPphB suppresses AvrB-induced activation of RPM1 but not AvrRpm1-induced activation. Molecular Plant-Microbe Interactions, 28(6), 727–735.

Santos, D., Martins da Silva, P., Abrantes, I., & Maleita, C. (2020). Tomato Mi-1.2 gene confers resistance to Meloidogyne luci and M. ethiopica. European Journal of Plant Pathology, 156(2), 571–580.

Sarris, P. F., Duxbury, Z., Huh, S. U., Ma, Y., Segonzac, C., Sklenar, J., Derbyshire, P., Cevik, V., Rallapalli, G., & Saucet, S. B. (2015). A plant immune receptor detects pathogen effectors that target WRKY transcription factors. Cell, 161(5), 1089–1100.

Saucet, S. B., Ma, Y., Sarris, P. F., Furzer, O. J., Sohn, K. H., & Jones, J. D. (2015). Two linked pairs of Arabidopsis TNL resistance genes independently confer recognition of bacterial effector AvrRps4. Nature communications, 6(1), 1–12.

Saur, I. M., Bauer, S., Kracher, B., Lu, X., Franzeskakis, L., Müller, M. C., Sabelleck, B., Kümmel, F., Panstruga, R., & Maekawa, T. (2019). Multiple pairs of allelic MLA immune receptor-powdery mildew AVRA effectors argue for a direct recognition mechanism. Elife, 8.

Saur, I. M. L., Panstruga, R., & Schulze-Lefert, P. (2021). NOD-like receptor-mediated plant immunity: from structure to cell death. Nat Rev Immunol, 21(5), 305–318. https://doi.org/10.1038/s41577-020-00473-z

Schnable, P. S., Ware, D., Fulton, R. S., Stein, J. C., Wei, F., Pasternak, S., Liang, C., Zhang, J., Fulton, L., Graves, T. A., Minx, P., Reily, A. D., Courtney, L., Kruchowski, S. S., Tomlinson, C., Strong, C., Delehaunty, K., Fronick, C., Courtney, B.,. .. Wilson, R. K. (2009). The B73 Maize Genome: Complexity, Diversity, and Dynamics. Science, 326(5956), 1112–1115. https://doi.org/doi:10.1126/science.1178534

Schneider, D., Saraiva, A., Azzoni, A., Miranda, H., de Toledo, M., Pelloso, A., & Souza, A. (2010). Overexpression and purification of PWL2D, a mutant of the effector protein PWL2 from Magnaporthe grisea. Protein expression and purification, 74(1), 24–31.

Schultink, A., Qi, T., Bally, J., & Staskawicz, B. (2019). Using forward genetics in Nicotiana benthamiana to uncover the immune signaling pathway mediating recognition of the Xanthomonas perforans effector XopJ4. New Phytologist, 221(2), 1001–1009.

Schwessinger, B., & Rathjen, J. P. (2017). Extraction of high molecular weight DNA from fungal rust spores for long read sequencing. In Wheat rust diseases (pp. 49–57). Springer.

Seeholzer, S., Tsuchimatsu, T., Jordan, T., Bieri, S., Pajonk, S., Yang, W., Jahoor, A., Shimizu, K. K., Keller, B., & Schulze-Lefert, P. (2010). Diversity at the Mla powdery mildew resistance locus from cultivated barley reveals sites of positive selection. Molecular plant-microbe interactions, 23(4), 497–509.

Seong, K., & Krasileva, K. (2022). Comparative computational structural genomics highlights divergent evolution of fungal effectors. bioRxiv.

Seto, D., Koulena, N., Lo, T., Menna, A., Guttman, D. S., & Desveaux, D. (2017). Expanded type III effector recognition by the ZAR1 NLR protein using ZED1-related kinases. Nature Plants, 3(4), 1–4.

Shao, F., Golstein, C., Ade, J., Stoutemyer, M., Dixon, J. E., & Innes, R. W. (2003). Cleavage of Arabidopsis PBS1 by a bacterial type III effector. Science, 301(5637), 1230–1233.

Shen, Q.-H., Zhou, F., Bieri, S., Haizel, T., Shirasu, K., & Schulze-Lefert, P. (2003). Recognition specificity and RAR1/SGT1 dependence in barley Mla disease resistance genes to the powdery mildew fungus. The Plant Cell, 15(3), 732–744.

Sirisathaworn, T., Srirat, T., Longya, A., & Jantasuriyarat, C. (2017). Evaluation of mating type distribution and genetic diversity of three Magnaporthe oryzae avirulence genes, PWL-2, AVR-Pii and Avr-Piz-t, in Thailand rice blast isolates. Agriculture and Natural Resources, 51(1), 7–14.

Smedley, M. A., & Harwood, W. A. (2015). Gateway®-compatible plant transformation vectors. In Agrobacterium Protocols (pp. 3–16). Springer.

Stewart, C. N., & Via, L. E. (1993). A rapid CTAB DNA isolation technique useful for RAPD fingerprinting and other PCR applications. Biotechniques, 14(5), 748–751.

Sun, J., Huang, G., Fan, F., Wang, S., Zhang, Y., Han, Y., Zou, Y., & Lu, D. (2017). Comparative study of Arabidopsis PBS1 and a wheat PBS1 homolog helps understand the mechanism of PBS1 functioning in innate immunity. Scientific reports, 7(1), 1–12.

Sweat, T. A., Lorang, J. M., Bakker, E. G., & Wolpert, T. J. (2008). Characterization of natural and induced variation in the LOV1 gene, a CC-NB-LRR gene conferring victorin sensitivity and disease susceptibility in Arabidopsis. Molecular plant-microbe interactions, 21(1), 7–19.

Sweat, T. A., & Wolpert, T. J. (2007). Thioredoxin h 5 is required for victorin sensitivity mediated by a CC-NBS-LRR gene in Arabidopsis. The Plant Cell, 19(2), 673–687.

Sweigard, J. A., Carroll, A. M., Kang, S., Farrall, L., Chumley, F. G., & Valent, B. (1995). Identification, cloning, and characterization of PWL2, a gene for host species specificity in the rice blast fungus. The plant cell, 7(8), 1221–1233.

Takahashi, H., Miller, J., Nozaki, Y., Sukamto, Takeda, M., Shah, J., Hase, S., Ikegami, M., Ehara, Y., & Dinesh-Kumar, S. (2002). RCY1, an Arabidopsis thaliana RPP8/HRT family resistance gene, conferring resistance to cucumber mosaic virus requires salicylic acid, ethylene and a novel signal transduction mechanism. The Plant Journal, 32(5), 655–667.

Talbot, N. J., Ebbole, D. J., & Hamer, J. E. (1993). Identification and characterization of MPG1, a gene involved in pathogenicity from the rice blast fungus Magnaporthe grisea. The Plant Cell, 5(11), 1575–1590.

Thind, A. K., Wicker, T., Šimková, H., Fossati, D., Moullet, O., Brabant, C., Vrána, J., Doležel, J., & Krattinger, S. G. (2017). Rapid cloning of genes in hexaploid wheat using cultivar-specific long-range chromosome assembly. Nature Biotechnology, 35(8), 793–796.

Toruño, T. Y., Shen, M., Coaker, G., & Mackey, D. (2019). Regulated disorder: posttranslational modifications control the RIN4 plant immune signaling hub. Molecular Plant-Microbe Interactions, 32(1), 56–64.

Tosa, Y., Osue, J., Eto, Y., Oh, H.-S., Nakayashiki, H., Mayama, S., & Leong, S. A. (2005). Evolution of an avirulence gene, AVR1-CO39, concomitant with the evolution and differentiation of Magnaporthe oryzae. Molecular Plant-Microbe Interactions, 18(11), 1148–1160.

Valent, B., Crawford, M. S., Weaver, C. G., & Chumley, F. G. (1986). Genetic studies of fertility and pathogenicity in Magnaporthe grisea(Pyricularia oryzae). Iowa State J. Res., 60(4), 569–594.

Van Der Biezen, E. A., & Jones, J. D. (1998). Plant disease-resistance proteins and the gene-for-gene concept. Trends in biochemical sciences, 23(12), 454–456.

Van der Hoorn, R. A., De Wit, P. J., & Joosten, M. H. (2002). Balancing selection favors guarding resistance proteins. Trends in plant science, 7(2), 67–71.

Van Ooijen, J. (2006). JoinMap 4. Software for the calculation of genetic linkage maps in experimental populations. Kyazma BV, Wageningen, Netherlands, 33.

Vogel, J. P., Garvin, D. F., Mockler, T. C., Schmutz, J., Rokhsar, D., Bevan, M. W., Barry, K., Lucas, S., Harmon-Smith, M., Lail, K., Tice, H., Schmutz, J., Grimwood, J., McKenzie, N., Bevan, M. W., Huo, N., Gu, Y. Q., Lazo, G. R., Anderson, O. D.,. .. gene family, a. (2010). Genome sequencing and analysis of the model grass Brachypodium distachyon. Nature, 463(7282), 763–768. https://doi.org/10.1038/nature08747

Vos, P., Simons, G., Jesse, T., Wijbrandi, J., Heinen, L., Hogers, R., Frijters, A., Groenendijk, J., Diergaarde, P., Reijans, M., Fierens-Onstenk, J., Both, M. d., Peleman, J., Liharska, T., Hontelez, J., & Zabeau, M. (1998). The tomato Mi-1 gene confers resistance to both root-knot nematodes and potato aphids. Nature Biotechnology, 16(13), 1365–1369. https://doi.org/10.1038/4350

Walker, B. J., Abeel, T., Shea, T., Priest, M., Abouelliel, A., Sakthikumar, S., Cuomo, C. A., Zeng, Q., Wortman, J., Young, S. K., & Earl, A. M. (2014). Pilon: an integrated tool for comprehensive microbial variant detection and genome assembly improvement. PLoS One, 9(11), e112963. https://doi.org/10.1371/journal.pone.0112963

Wang, G., Roux, B., Feng, F., Guy, E., Li, L., Li, N., Zhang, X., Lautier, M., Jardinaud, M.-F., & Chabannes, M. (2015). The decoy substrate of a pathogen effector and a pseudokinase specify pathogen-induced modified-self recognition and immunity in plants. Cell host & microbe, 18(3), 285–295.

Wang, X., Richards, J., Gross, T., Druka, A., Kleinhofs, A., Steffenson, B., Acevedo, M., & Brueggeman, R. (2013). The rpg4-mediated resistance to wheat stem rust (Puccinia graminis) in barley (Hordeum vulgare) requires Rpg5, a second NBS-LRR gene, and an actin depolymerization factor. Molecular plant-microbe interactions, 26(4), 407–418.

Waterhouse, R. M., Seppey, M., Simão, F. A., Manni, M., Ioannidis, P., Klioutchnikov, G., Kriventseva, E. V., & Zdobnov, E. M. (2018). BUSCO Applications from Quality Assessments to Gene Prediction and Phylogenomics. Mol Biol Evol, 35(3), 543–548. https://doi.org/10.1093/molbev/msx319

Wei, F., Gobelman-Werner, K., Morroll, S. M., Kurth, J., Mao, L., Wing, R., Leister, D., Schulze-Lefert, P., & Wise, R. P. (1999). The Mla (powdery mildew) resistance cluster is associated with three NBS-LRR gene families and suppressed recombination within a 240-kb DNA interval on chromosome 5S (1HS) of barley. Genetics, 153(4), 1929–1948.

Wei, F., Wing, R. A., & Wise, R. P. (2002). Genome dynamics and evolution of the Mla (powdery mildew) resistance locus in barley. The Plant Cell, 14(8), 1903–1917.

Williams, S. J., Sohn, K. H., Wan, L., Bernoux, M., Sarris, P. F., Segonzac, C., Ve, T., Ma, Y., Saucet, S. B., & Ericsson, D. J. (2014). Structural basis for assembly and function of a heterodimeric plant immune receptor. Science, 344(6181), 299–303.

Witek, K., Jupe, F., Witek, A. I., Baker, D., Clark, M. D., & Jones, J. D. (2016). Accelerated cloning of a potato late blight–resistance gene using RenSeq and SMRT sequencing. Nature biotechnology, 34(6), 656–660.

Wu, C.-H., Abd-El-Haliem, A., Bozkurt, T. O., Belhaj, K., Terauchi, R., Vossen, J. H., & Kamoun, S. (2017). NLR network mediates immunity to diverse plant pathogens. Proceedings of the National Academy of Sciences, 114(30), 8113–8118. https://doi.org/10.1073/pnas.1702041114

Zhou, F., Kurth, J., Wei, F., Elliott, C., Valè, G., Yahiaoui, N., Keller, B., Somerville, S., Wise, R., & Schulze-Lefert, P. (2001). Cell-autonomous expression of barley Mla1 confers race-specific resistance to the powdery mildew fungus via a Rar1-independent signaling pathway. The Plant Cell, 13(2), 337–350.

Zimin, A. V., Marçais, G., Puiu, D., Roberts, M., Salzberg, S. L., & Yorke, J. A. (2013). The MaSuRCA genome assembler. Bioinformatics, 29(21), 2669–2677.

Zuker, M. (2003). Mfold web server for nucleic acid folding and hybridization prediction. Nucleic acids research, 31(13), 3406–3415.

Zwyrtková, J., Šimková, H., Doležel, J. (2021). Chromosome genomics uncovers plant genome organization and function. Biotechnology Advances 46, 107659.

